# CLCC1 Governs Bilayer Equilibration at the Endoplasmic Reticulum to Maintain Cellular and Systemic Lipid Homeostasis

**DOI:** 10.1101/2024.06.07.596575

**Authors:** Lingzhi Wu, Jianqin Wang, Yawei Wang, Junhan Yang, Yuanhang Yao, Dong Huang, Yating Hu, Xinxuan Xu, Renqian Wang, Wenjing Du, Yiting Shi, Quan Li, Lu Liu, Yuangang Zhu, Xiao Wang, Qiang Guo, Li Xu, Peng Li, Xiao-Wei Chen

## Abstract

The intricate orchestration of lipid production, storage, and mobilization is vital for cellular and systemic homeostasis^1,2^. Dysfunctional plasma lipid control represents the major risk factor for cardio-metabolic diseases, the leading cause of human mortality^3,4^. Within the cellular landscape, the endoplasmic reticulum (ER) is the central hub of lipid synthesis and secretion, particularly in metabolically active hepatocytes in the liver or enterocytes in the gut^5,6^. Initially assembled in the ER lumen, lipid-ferrying lipoproteins necessitate the cross-membrane transfer of both neutral and phospho-lipids onto the lumenal apolipoprotein B (APOB), in a poorly-defined process^7–10^. Here we show that trans-bilayer equilibration of phospholipids, regulated by the ER protein CLCC1, determines lipid partition across the ER membrane and consequently systemic lipid homeostasis. CLCC1 partners with the phospholipid scramblase TMEM41B^11,12^ to recognize imbalanced bilayers and promote lipid scrambling, thereby licensing lipoprotein biogenesis and the subsequent bulk lipid transport. Strikingly, loss of CLCC1 or TMEM41B leads to the emergence of giant lumenal lipid droplets enclosed by extensively imbalanced ER bilayers, and consequently drastically accelerated pathogenesis of metabolic-dysfunction-associated liver steatohepatitis (MASH). The above results establish phospholipid scrambling at the ER as the lynchpin to maintain a dynamic equilibrium. Considering the requirement of trans-bilayer phospholipid equilibration in numerous biological processes, ranging from catabolic autophagy to viral infection^13–16^, our study may enable further elucidation of a previously under-appreciated homeostatic control mechanism intrinsic to the ER function in lipid biogenesis and distribution.

## Leaflet imbalance directs lipids into <u>g</u>iant <u>E</u>R-enclosed lipid droplets (geLD)

The cross-bilayer transverse of amphipathic phospholipids synthesized asymmetrically on the cytosolic leaflet of the ER bilayer has remained an unaddressed problem until the recent identification of biogenic lipid scramblases, particularly TMEM41B^9,11,12^. Remarkably, deficient phospholipid scrambling due to hepatic TMEM41B inactivation not only blocks neutral lipid loading and the biogenesis of very-low-density lipoproteins (VLDLs), but also triggers massive lipid over-production and drastically accelerates metabolic-dysfunction-associated liver steatohepatitis (MASH)^11^. Unexpectedly, transmission electron microscopy (TEM) revealed that the innumerable lipid droplets in TMEM41B deficient hepatocytes were all tightly surrounded by membranes (Figure 1a and Extended Data Figure 1a), likely originated from the ER. To further understand these unusual lipid storages, we employed the newly-developed electron tomography coupled with high pressure cryo-fixation (cryo-ET) of hepatic tissues^17,18^. As expected^19,20^, lipid droplets in wild type hepatocytes were observed to be situated in the cytosol (Figure 1b, left), often juxtaposed to the ER with an average lumen width of ∼50 nm (Figure 1b, arrowhead). By contrast, lipid droplets in TMEM41B-deficient hepatocytes were enclosed within the ER bilayer and exhibited an average diameter of ∼1μm (Figure 1b, right). Subsequent segmentation and 3D reconstruction revealed that these giant lipid droplets were tightly wrapped within the ER lumen by a single ER bilayer characterized by the smooth, ribosome-free appearance, while displaying markedly increased curvature (Figure 1c, Extended Data Figure 1b and Extended Data Video 1&2). Collectively, the ultrastructural analysis uncovered an unusual lipid storage, featured by the emergence of giant ER-enclosed lipid droplets (geLD) within the ER lumen, accompanied by highly curved ER bilayers with imbalanced leaflets.

**Figure 1.**
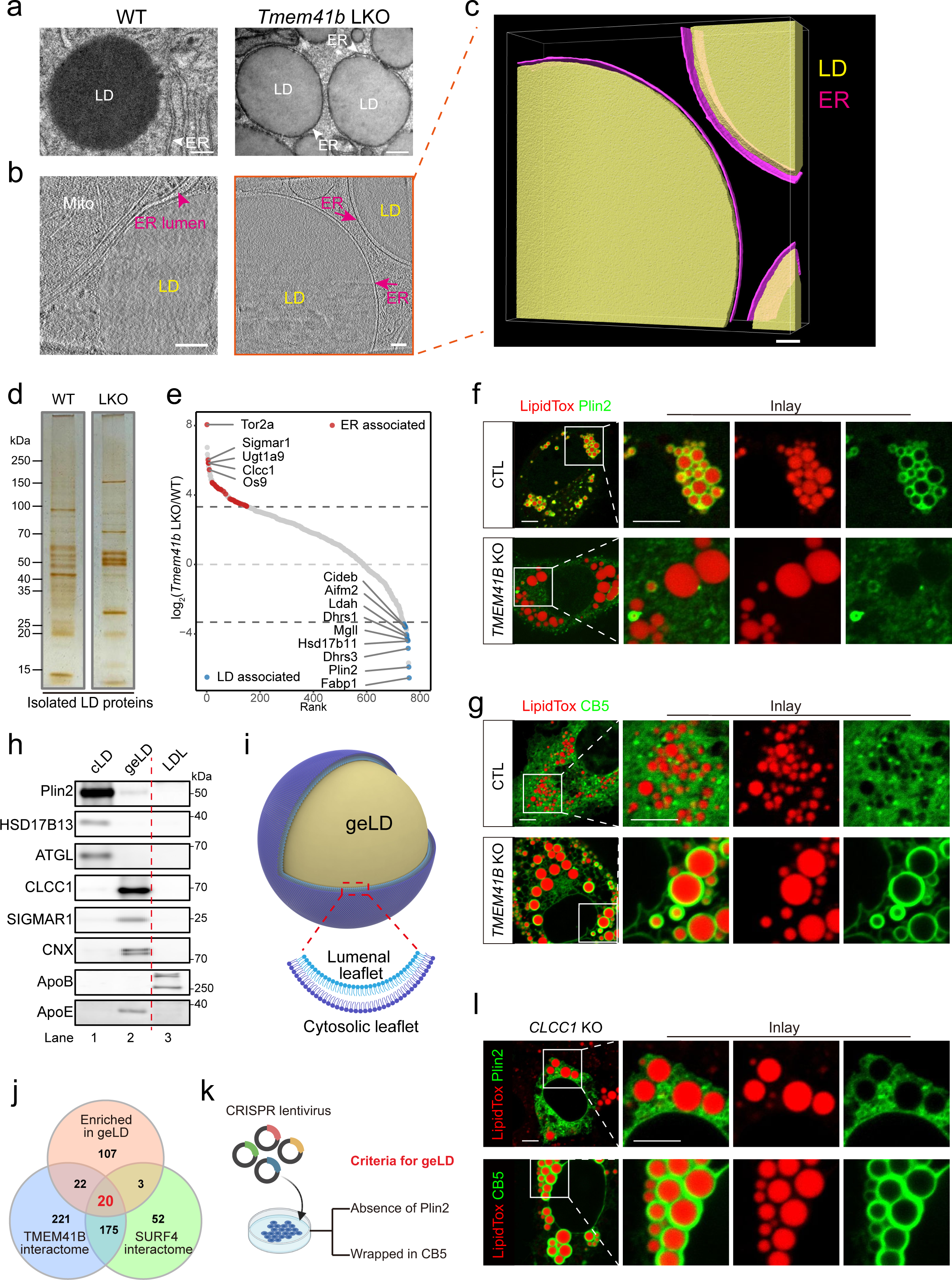
Leaflet imbalance directs lipids into <u>g</u>iant <u>E</u>R-enclosed lipid droplets (geLD) **a.** Representative transmission electron microscopy of livers from wild type (WT, left) or *Tmem41b* liver-specific knockout (LKO, right) mice. n = 5 mice for each genotype. Arrowheads indicate the ER in WT and *Tmem41b* LKO hepatocytes. LD, lipid droplets. Scale bars, 200 nm. **b.** Electron tomography of high pressure cryo-fixed (cryo-ET) liver samples from WT (left) or *Tmem41b* LKO (right) mice. Representatives of 114 views from 2 WT mice livers and 177 views from 2 *Tmem41b* LKO mice livers are shown. Arrowhead indicates the ER lumen in WT hepatocytes (left) and arrows indicate the ER bilayer in *Tmem41b* KO hepatocytes (right). Mito, mitochondria. Scale bars, 100 nm. **c.** Three-dimensional rendering of tomogram corresponding to the region of *Tmem41b* KO hepatocytes in (b). Yellow: LD, magenta: ER membrane. Scale bars, 100 nm. **d.** Protein composition of LDs isolated from WT (left) and *Tmem41b* LKO (right) liver. Same quantity of proteins from both samples were separated by SDS-PAGE and visualized by sliver staining. **e.** Quantitative proteomics of hepatic LD isolated from WT and *Tmem41b* LKO mice. Significantly altered ER-associated (red) or cytosolic LD-associated (blue) proteins are listed. **f.** Differential label of cLD and geLD by the cLD marker Plin2. Confocal microscopy of control (upper) and *TMEM41B* KO (lower) Huh7 cells infected with AAVs expressing GFP-Plin2. Red, LipidTOX. Scale bar, 5 μm. **g.** Differential label of cLD and geLD by the ER marker GFP-CB5. Confocal microscopy of the same sets of cells as in (f) infected with AAVs expressing GFP-CB5 (amino acids 90 to 124 of Cytochrome B5 type A). Red, LipidTOX. Scale bar, 5 μm. **h.** IB of cLDs isolated from WT (lane1), geLDs from *Tmem41b* LKO (lane2) mice liver and plasma LDL (lane3). LDL, low-density-lipoproteins. Similar quantity of proteins from each sample were loaded onto the gels. **i.** Model depicting a geLD that is accompanied by extensively curved ER bilayers in TMEM41B-deficient hepatocytes. **j.** Venn diagram of geLD-enriched proteome, TMEM41B interactome, and SURF4 interactome detected by mass spectrometry. **k.** Schematic diagram of the targeted CRISPR/Cas9 screen for geLDs. **l.** CLCC1 deficiency induces geLDs. CRISPR/Cas9-mediated *CLCC1* KO Huh7 cells were infected with AAVs expressing GFP-Plin2 (upper) or GFP-CB5 (lower), and subsequently stained with LipidTOX for confocal microscopy analysis. Scale bar, 5 μm. For **d-h** and **l**, representative results of at least 3 biological independent replicates are shown.

Biochemical isolation and silver staining revealed distinctive protein compositions in conventional cytosolic lipid droplets (cLDs) purified from wild type liver and geLDs from TMEM41B-deficient liver (Figure 1d and Extended Data Figure 1c). Subsequent quantitative mass spectrometry revealed that marker proteins associated with cLD, notably Perilipin 2 (Plin2)^21,22^, were depleted from geLD (Figure 1e, blue dots). Instead, these unusual lipid droplets exhibited enrichment of ER membrane proteins, such as Torsins, Sigma-1 Receptor, and Ugt1a9 (Figure 1e, red dots and Extended Data Figure 1d). Consistent with these mass spectrometry data, confocal microscopy confirmed that Plin2 delineated the surface of cLD in wild type cells, but was devoid from geLDs in TMEM41B-deficient cells (Figure 1f). Conversely, the geLDs were encircled by an ER membrane marker generated from the cytochrome B5 protein (GFP-CB5), which was absent from cLD in wild type cells (Figure 1g). Immuno-blotting (IB) further confirmed the absence of cLD markers including Plin2, HSD17B13, or the lipolytic enzyme ATGL from geLDs and the enrichment of a subset of ER membrane proteins (Figure 1h, lane 1 vs 2). Of note, geLDs were also devoid of apolipoprotein B (APOB), the principal structural protein of VLDLs (Figure 1h, lane 2 vs 3). By contrast, geLDs contained the exchangeable apolipoprotein APOE^23^.

The combination of cryo-ET, microscopy, and biochemical characterizations collectively unveiled the emergence of the unique geLD storage, accompanied by extensively curved ER bilayers and consequently sustained imbalance between the cytosolic vs the lumenal leaflets (Figure 1i). The emergence of massive imbalanced leaflets prompted us to hypothesize that geLDs may in turn enrich factors promoting ER bilayer equilibration, coincided with efficient phospholipid scrambling and also lipoprotein secretion. To test this hypothesis, we compiled the geLD enriched proteome with the interactomes associated with the TMEM41B scramblase^11^ and the SURF4 cargo receptor^24^ (Figure 1j), resulting in a short list of proteins led by a poorly understood ER membrane protein CLCC1 (chloride channel CLIC like 1). Subsequently, we initiated a targeted CRISPR-mediated screen focusing on the hits identified in figure 1j, guided by the characteristics of geLD (Figure 1k). Of note, loss of CLCC1 led to the induction of geLDs featured by the absence of Plin2 decoration but instead encircled with GFP-CB5 (Figure 1l), recapitulating those resulted from TMEM41B deficiency. Of note, the rough ER marker GFP-SEC61β showed little enrichment around geLDs in TMEM41B or CLCC1 deficient cells and the ER lumen marker GFP-KEDL exhibited moderate enrichment (Extended Data Figure 2a&b), whereas ApoE-GFP displayed marked enrichment to delineate the surface of geLDs (Extended Data Figure 2c), implicating these unique structures in lipid regulation.

## CLCC1 partners with TMEM41B scramblase in governing bulk lipid secretion

Consistent with their convergence of cellular function uncovered above, tandem affinity purification revealed biochemical associations between CLCC1 and TMEM41B (Figure 2a). Accordingly, endogenous CLCC1 could be isolated from the endogenous TMEM41B immune-complex (Figure 2b). Moreover, genetic analysis based on the recently available GLGC (Global Lipid Genetics Consortium) data^25^ uncovered strong associations between genetic variants in the human *CLCC1* gene and plasma lipids in populations (Figure 2c&d and Extended Data Figure 3), particularly in atherogenic LDL levels (p = 1.69E^-41^, β = −0.078). Of note, the minor allele is present in rather low frequency in different ethnicities (less than or close to 0.01, Figure 2e), indicating possible strong biological effects of CLCC1 on lipid homeostasis.

**Figure 2.**
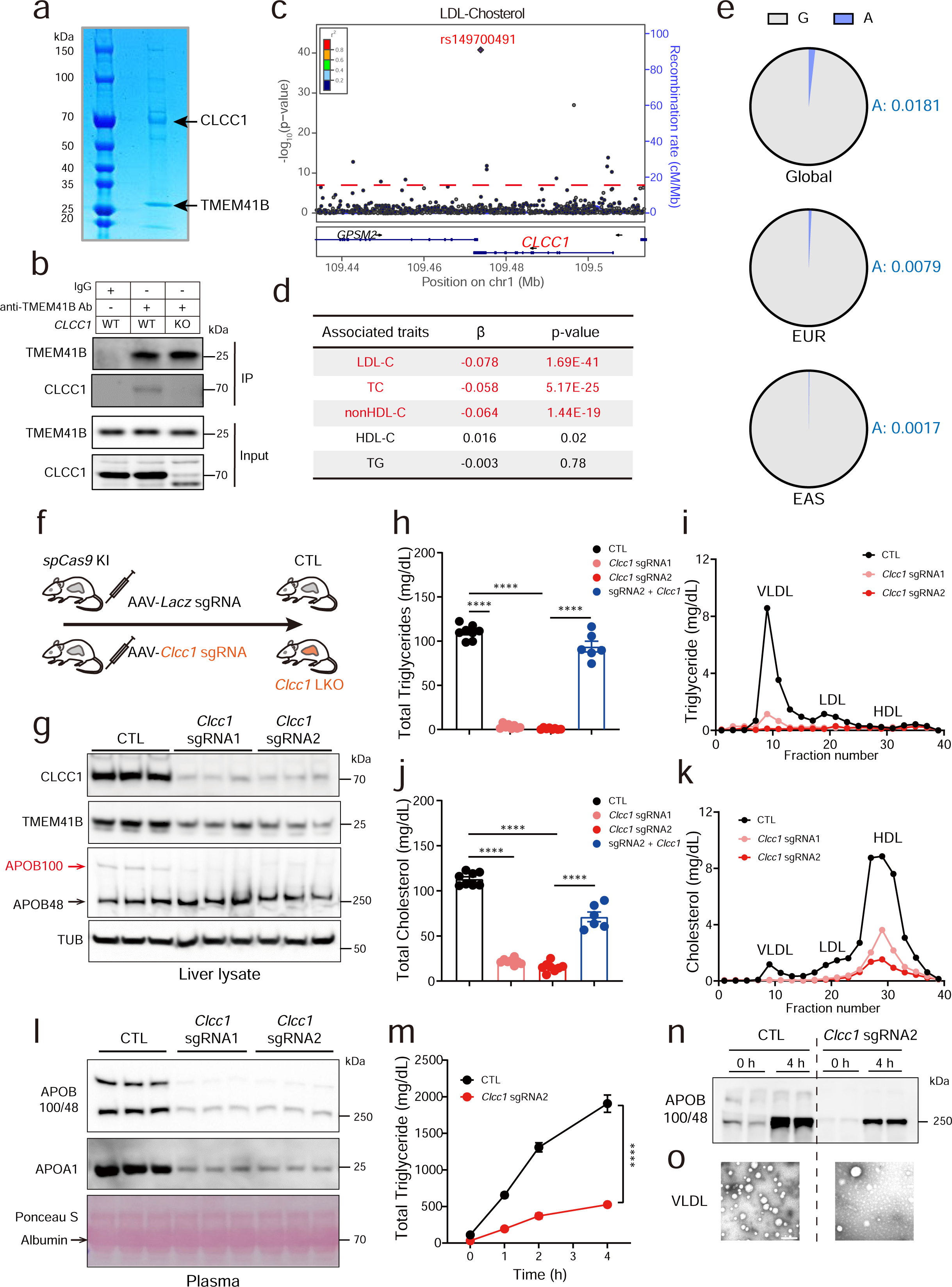
CLCC1 partners with TMEM41B scramblase in governing bulk lipid secretion. **a.** Coomassie blue staining of the CLCC1-TMEM41B complex. Arrows indicate CLCC1 (upper) and TMEM41B (lower), respectively. HEK293F cells expressing CLCC1-FLAG and twin-StrepTagII-TMEM41B were lysed and subjected to tandem purification sequentially with anti-FLAG resins and Streptactin affinity beads. **b.** Co-immunoprecipitation (co-IP) of endogenous TMEM41B and CLCC1 in Huh7 cells. Lysates from cells with indicated genotypes were subjected to anti-TMEM41B IP or anti-control IgG IP, prior to SDS-PAGE and IB with the indicated antibodies. **c.** Regional plot of *CLCC1* associated with plasma LDL-cholesterol levels in humans. **d.** Summary of the GLGC genome-wide association data between the leading SNP rs149700491 in *CLCC1* and plasma lipids in humans. **e.** Minor allele frequency (MAF) difference at SNP rs149700491 in European (EUR) and East Asian (EAS) descents from the 1000 Genome Project (Global). **f.** Schematic diagram of CRISPR/Cas9-mediated acute hepatic gene inactivation. Top: *Lacz* targeting sgRNA as control (CTL), bottom: *Clcc1* targeting sgRNA. **g.** Hepatic CLCC1 inactivation leads to depletion of TMEM41B and APOB100. Total liver protein from mice receiving *Lacz* sgRNA control (CTL) or 2 different *Clcc1* targeting sgRNA were conducted with IB analysis. **h.** Circulating triglyceride (TG) levels in control (CTL) or *Clcc1* LKO mice. Plasma of *Lacz* sgRNA control (n = 8), *Clcc1* sgRNA1 (n = 8), *Clcc1* sgRNA2 (n = 8) and *Clcc1* sgRNA2 plus CRISPR-resistant *Clcc1* cDNA (n = 6) were collected and analyzed. Data are presented as mean ± SEM. ****p < 0.0001 (two-tailed Student’s t test). **i.** Plasma samples from CTL or *Clcc1* LKO mice in (h) were fractionated by fast-protein liquid chromatography (FPLC) followed by TGs measurements. VLDL, very-low-density lipoproteins. LDL, low-density lipoproteins. HDL, high-density lipoproteins. **j.** Circulating cholesterol levels in (h). Data are presented as mean ± SEM. ****p < 0.0001 (two-tailed Student’s t test). **k.** Cholesterol measurement of samples in (i). **l.** IB analysis of plasma from CTL or *Clcc1* LKO mice. **m.** Hepatic VLDL secretion from CTL or *Clcc1* LKO mice. Plasma samples were collected from *Lacz* sgRNA control (CTL) (n = 4) and *Clcc1* sgRNA2 (n = 4) mice injected with the LPL inhibitor (Tyloxapol), and plasma TG concentrations over time were determined as a readout of VLDL secretion. Data are presented as mean ± SEM. ****p<0.0001 (two-tailed Student’s t test). **n.** IB analysis of plasma APOB from (m). Each sample contains plasma pooled from two mice. **o.** Negative staining of VLDL particles from (j-k). Scale bar, 200 nm. For **a-b, g, i, k-l**, and **n-o**, representative results of at least 3 biological independent replicates are shown.

The biochemical and genetic data prompted us to investigate the physiological function of CLCC1, focusing on lipoprotein biogenesis and transport in which TMEM41B plays an essential role. We thus employed CRISPR-mediated *in vivo* gene editing^26^ to acutely inactivate hepatic *CLCC1* in murine models (Figure 2f and Extended Data Figure 4a). Interestingly, IB analysis of liver proteins showed that CLCC1 depletion by two different sgRNAs both led to the down-regulation of TMEM41B protein without altering its transcript levels (Figure 2g and Extended Data Figure 4b), further indicating physical associations between these two transmembrane proteins. Moreover, APOB100 protein also became depleted, a phenomenon known to reflect the failure in APOB lipidation and therefore in the subsequent VLDL biogenesis^27–29^. Of note, such defects with APOB100 were also observed with hepatic TMEM41B inactivation^11^, together pointing to an essential role of the trans-bilayer phospholipid supply in the initiating step of VLDL biogenesis.

Strikingly, CRISPR-mediated inactivation of hepatic CLCC1 led to depletion of plasma triglycerides to near zero in fasted mice (Figure 2h, column 1 vs 2&3). This drastic lipid depletion could be rescued by re-introduction of sgRNA-resistant CLCC1 (Figure 2h, column 4 and Extended Data Figure 4c), demonstrating the specificity of the lipid lowering effects to CLCC1 deficiency. Profiling of plasma lipids via size exclusion chromatography also confirmed the depletion of atherogenic lipoproteins including VLDL and LDL (Figure 2i). Similar reduction in plasma cholesterol were also observed (Figure 2j&k). Consistent with the lipid measurements, circulating apolipoproteins including APOB and APOA1 were also depleted in the plasma of hepatic CLCC1-deficiency mice, while Albumin levels remained unaltered (Figure 2l). Accordingly, the mutant mice exhibited significant reduction in hepatic triglyceride secretion compared to controls upon injection of the LPL inhibitor Tyloxapol^24^ (Figure 2m). Examination of circulating APOB protein levels or visualization of isolated lipoproteins by EM negative staining further confirmed diminished lipoprotein secretion caused by hepatic CLCC1 inactivation (Figure 2n&o).

## CLCC1 recognizes imbalanced leaflets and promotes phospholipid scrambling

The essential role of CLCC1 in lipoprotein-mediated lipid homeostasis intrigued us to determine its underlying molecular mechanism in lipid control. Interestingly, alphafold3-assisted prediction^30^ showed that CLCC1 may exist in high-molecular weight oligomeric forms that assemble into a ring-like configuration (Figure 3a&b). We then performed blue native PAGE to analyze CLCC1, and observed an induction of CLCC1 oligomers in TMEM41B-deficient cells (Figure 3c). Notably, CLCC1 oligomers were enriched in ER membranes encircling geLDs (Figure 3c, right two lanes), indicating a preferred localization to the imbalanced ER bilayers induced by scramblase deficiency. Consistent with this notion, confocal microscopy revealed that endogenous CLCC1 exhibited a diffusive localization throughout the ER in wild type cells (Figure 3d, upper panels), whereas in TMEM41B-deficient cells, CLCC1 was recruited and enriched on the curved ER membranes delineating geLDs (Figure 3d, lower panels). By contrast, TMEM41B failed to localize to the imbalanced ER bilayers in CLCC1-deficient cells (Extended Data Figure 5a).

**Figure 3.**
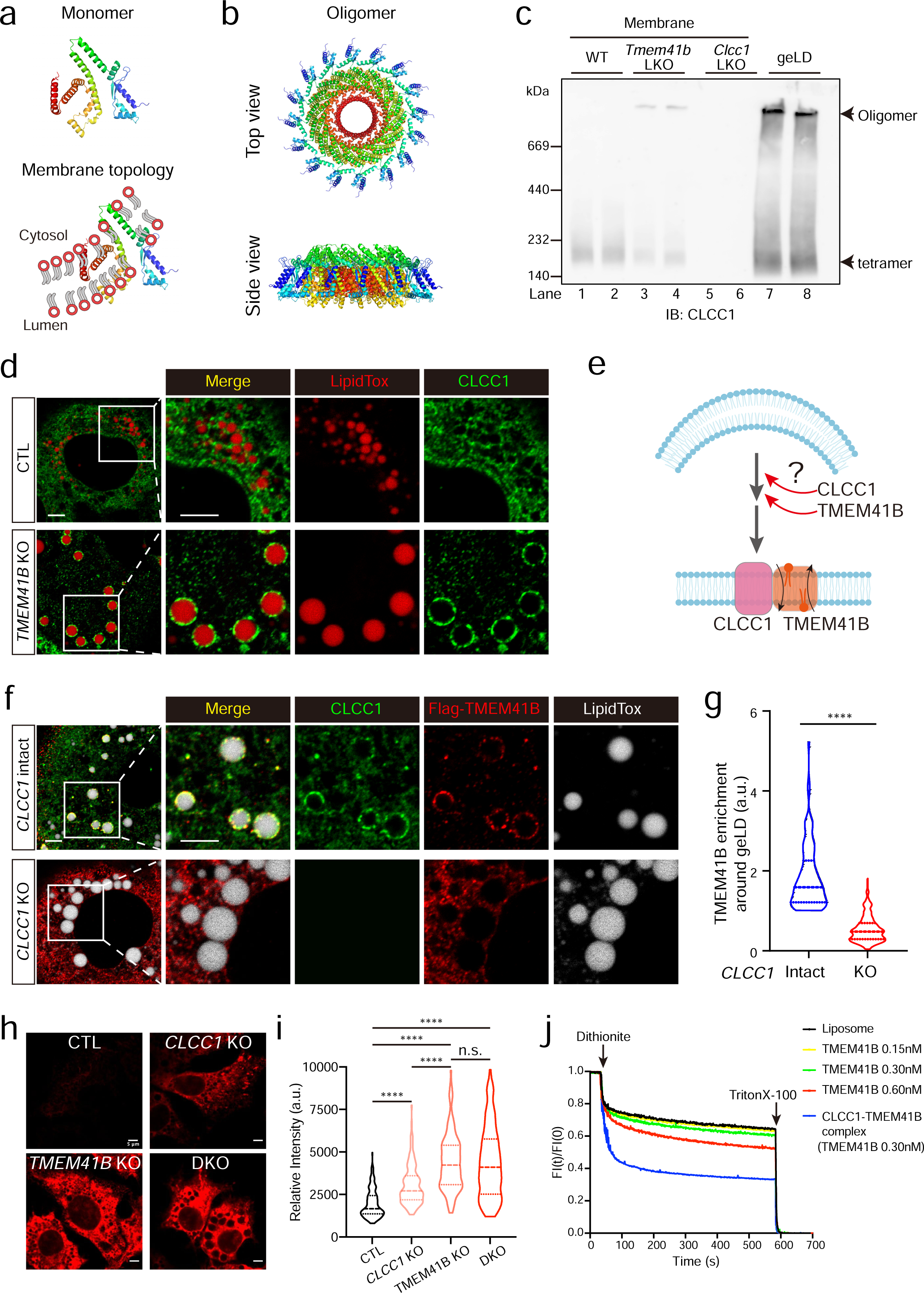
CLCC1 promotes TMEM41B scramblase-catalyzed phospholipid equilibration *in vivo* and *in vitro*. **a.** AlphaFold3-predicted structure of human CLCC1 monomer (amino acids 90 to 360). Blue to red represents the N-to C-terminal orientation of CLCC1. **b.** AlphaFold3-predicted hexadecameric structure of human CLCC1 form (a). Top view (upper) refers to the perspective from the cytosol towards the lumen. **c.** CLCC1 oligomerization in TMEM41B-deficient cell. Blue native PAGE analysis of CLCC1 in membranes isolated from WT (lane 1, 2), *Tmem41b* LKO (lane 3, 4), *Clcc1* LKO (lane 5, 6) liver and geLDs (lane 7, 8) from *Tmem41b* LKO liver. **d.** CLCC1 recognizes ER bilayers encircling geLDs. Control (upper) and *TMEM41B* KO (lower) Huh7 cells were co-stained with an anti-CLCC1 antibody and LipidTOX prior to confocal microscopy. Scale bar, 5 μm. **e.** Working model depicting the CLCC1-dependent TMEM41B recruitment to imbalanced leaflets to catalyze bilayer equilibration. **f.** CLCC1-dependent recruitment of TMEM41B to the ER membranes encircling geLD. Control (upper) and *CLCC1* KO (lower) generated in TMEM41B-deficient Huh7 cells were infected with AAVs expressing FLAG-TMEM41B, and were co-stained with anti-CLCC1, anti-FLAG antibodies and LipidTOX prior to confocal microscopy. Scale bar, 5 μm. **g.** Quantitative analysis of geLD-associated FLAG TMEM41B signals from (f). Data are presented as violin plots. ****p<0.0001 (two-tailed Student’s t test). n = 100 and 210 LDs from 10 *CLCC1* intact and 14 *CLCC1* KO cells, respectively. **h.** CLCC1 inactivation impairs lipid scrambling at the ER. CRISPR/Cas9-mediated control, *CLCC1* KO, *TMEM41B* KO and *CLCC1*/*TMEM41B* double-KO Huh7 cells were incubated with alkyne-choline, fixed, and permeabilized with digitonin to preserve the integrity of the ER bilayer. Newly synthesized phosphatidylcholines (PC) containing the alkyne group were visualized by “click reacted” with a fluorescent group to assess alkyne-PC levels at the outer side leaflet of ER, as a readout of lipid scrambling *in vivo*. **i.** Quantitative analysis of metabolically labeled PC signals in (h). n ≥ 13 cells from each group from (h). Data are presented as violin plots. ****p<0.0001 (two-tailed Student’s t test), n. s., not significant. **j.** CLCC1 promotes TMEM41B-mediated lipid scrambling *in vitro*. Liposomes contained NBD labelled PC were reconstituted with buffer alone, TMEM41B in different doses, and recombinant CLCC1-TMEM41B complex for *in vitro* lipids scrambling assay. Dithionite was added to quench NBD-PC signals at the outside leaflet of membranes, and Triton X-100 was added to permeabilize the membranes for complete fluorescence quenching. For **c**-**d, f, h** and **j**, representative results of at least 3 biological independent replicates are shown.

The above biochemical and localization data led us to test whether CLCC1 may be required for targeting the scramblase to imbalanced ER bilayers and subsequently equilibrate such asymmetry (Figure 3e). We thus transiently reconstituted FLAG-tagged TMEM41B into cells deficient of the scramblase. Indeed, FLAG-tagged TMEM41B was recruited to the curved membranes encircling geLDs in the presence of cellular CLCC1 (Figure 3f, upper panels). By contrast, such recruitment failed to take place upon CRISPR-mediated inactivation of CLCC1 (Figure 3f, lower panels, quantified in 3g).

To test the afore-described spatial regulation by CLCC1 may impact phospholipid equilibration at the ER membrane, we employed a cell-based chemical biology assay^31^, wherein defective *in vivo* phospholipid scrambling across the ER bilayer would result in an accumulation of alkyne-labeled phosphatidylcholine (PC) in cells (Extended Data Figure 5b). Loss of the TMEM41B scramblase in cells caused PC accumulation (Figure 3h), as expected. Interestingly, CLCC1 inactivation also led to PC accumulation, albeit to a lesser extent than TMEM41B. Combined deficiency of CLCC1 and TMEM41B didn’t further exacerbate scrambling defects caused by TMEM41B inactivation alone (quantified in Figure 3i), pointing to a regulatory role of CLCC1 in scramblase-mediated bilayer equilibration.

We further utilized the *in vitro* scrambling assay^32^ to quantitatively assess CLCC1’s contribution in the cross-bilayer shuttling of phospholipids, reflected by the quenching of fluorescently-labeled phospholipids distributed on both leaflets of liposomes (Extended Data Figure 6a). As expected, incorporation of recombinant TMEM41B scramblase at low, intermediate, or high concentration led to a dosage-dependent reduction of fluorescence in the assay (Figure 3i). Of note, while recombinant CLCC1 displayed no detectable scramblase activity *in vitro* (Extended Data Figure 6b), incorporation of an intermediate dose of TMEM41B in complex of CLCC1 displayed substantially increased scrambling activity (Figure 3i and Extended Data Figure 6c). In all conditions, no leakage of the proteo-liposome was detected (Extended Data Figure 6d&e), further demonstrating promoting effect of CLCC1 on lipid scrambling catalyzed by the TMEM41B scramblase. Taken together, the biochemical, cellular, and *in vitro* reconstitution experiments collectively demonstrated a pivotal role of CLCC1 in governing scramblase-mediated bilayer equilibration.

## CLCC1 maintains hepatic ER architecture and tissue homeostasis

The fundamental requirement of bilayer equilibration at the ER in biology prompted us to further investigate the contribution of CLCC1 to tissue homeostasis. Notably, CRISPR-mediated hepatic inactivation of CLCC1 resulted in drastically accelerated pathogenesis into metabolic dysfunctions associated with steatohepatitis (MASH) as early as in 4 weeks, without additional diet challenge. The mutant livers were characterized by pathologies including whitening appearance, hepatocyte ballooning, fibrosis, and immune cell infiltration (Figure 4a). Importantly, these pathologic defects could all be rescued by the re-introduction of CLCC1, demonstrating the specificity of the gene inactivation. Compared to controls, normalized weights of CLCC1-defecient liver became nearly doubled (Figure 4b). Strikingly, the mutant liver could float to the surface in water, contrasting the sinking of wild type or CLCC1-rescued mutant livers (Figure 4c). Accordingly, the mutant liver exhibited substantial accumulation of hepatic lipids (Figure 4d), accompanied with severe liver damages as reflected by elevated liver enzymes in the circulation compared to controls (Figure 4e and Extended Data Figure 7a). Consistent with the histological defects, these global and molecular pathologies could all be rescued by CLCC1 re-introduction, further demonstrating the essential role of CLCC1 in liver homeostasis.

**Figure 4.**
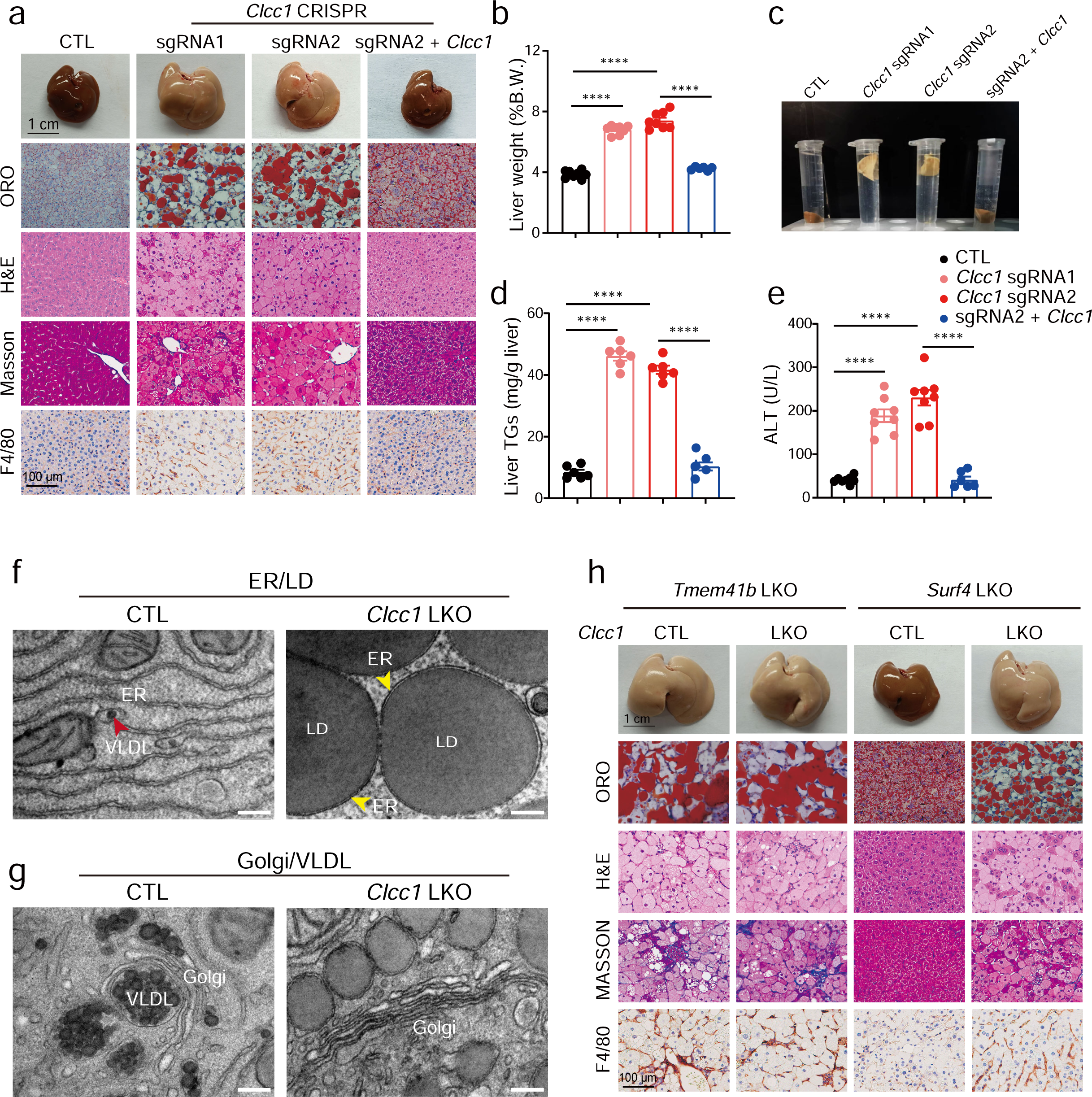
CLCC1 maintains hepatic ER architecture and tissue homeostasis. **a.** Hepatic CLCC1 inactivation leads to rapid progression into MASH. Livers from mice receiving indicated sgRNA or sgRNA plus CRSIPR-resistant *Clcc1* cDNA were subjected to the indicated pathology assessment. Mice were analyzed 4 weeks after AAVs delivery. Scale bars: upper (1 cm), lower (100 μm). Representative images from *Lacz* control (n = 8), *Clcc1* sgRNA1 (n = 8), *Clcc1* sgRNA2 (n = 8) or *Clcc1* sgRNA2 plus *Clcc1* (n = 6) are shown. **b.** Quantification of liver weight (% of body weight) of mice in (a). Data are presented as mean ± SEM. ****p < 0.0001 (two-tailed Student’s t test). **c.** Altered tissue density of CLCC1-deficient livers, which could float to the surface in water. **d.** Quantification of liver TG levels of mice in (a). Data are presented as mean ± SEM. ****p < 0.0001 (two-tailed Student’s t test). **e.** Quantification of plasma ALT of mice in (a). Data are presented as mean ± SEM. ****p < 0.0001 (two-tailed Student’s t test). **f.** TEM images of the hepatic LD and ER from WT (left) or *Clcc1* LKO (right) mice. Scale bars, 200 nm. Representative images of 5 independent mice for each group are shown. **g.** TEM images of the hepatic Golgi and VLDL from WT (left) or *Clcc1* LKO (right) mice. Representative images of 5 independent mice for each group are shown. **h.** Epistatic analysis of *Clcc1*/*Tmem41b* and *Surf4*. Liver samples from indicated mice with single or double gene deficiency were subjected to the indicated pathology assessments. Analysis was conducted 4 weeks post AAVs administration. Scale bars: upper (1 cm), lower (100 μm). Representative images of *Tmem41b* LKO (n = 7), *Clcc1/Tmem41b* LDKO (n = 7), *Surf4* LKO (n = 6) and *Surf4/Tmem41b* LDKO (n = 6) are shown.

We further performed ultrastructural analysis by TEM. As expected, the ER in wild type hepatocytes exhibited elaborate reticulum structure, characterized by regular lumen width and smooth ends often containing lipid particles of pre-VLDL size (<100 nm in diameter). By contrast, loss of CLCC1 led to the emergence of numerous geLDs (often >1μm in diameter), encircled by highly curved ER membrane (Figure 4f). Accordingly, VLDLs filled the Golgi apparatus in wild type hepatocytes, but were completely absent in the Golgi of CLCC1-deficient hepatocytes (Figure 4g). These data were consistent with the failure in APOB lipidation observed in Figure 2, further implicating the pivotal role of CLCC1 in phospholipid scrambling that licenses ApoB lipidation and VLDL assembly.

We lastly designed epistatic analysis to genetically pinpoint the function of CLCC1 in lipid regulation (Figure 4h). As previously reported, loss of hepatic TMEM41B in mice led to rapid emergence of MASH phenotypes, to a slightly greater extent than CLCC1 inactivation alone. However, combined deficiency of both TMEM41B and CLCC1 didn’t further exacerbate liver pathologies compared to the single deficiency of TMEM41B (Figure 4h, lane 1 vs 2). These genetic data strongly suggested that these two factors converge into the same step of lipid scrambling, thereby mediating leaflet equilibration and lipoprotein biogenesis. By contrast, blocking lipoprotein export by inactivation of the cargo receptor SURF4 didn’t cause overt liver pathogenesis. However, combined deficiency of SURF4 and CLCC1 produced phenotypes similar to CLCC1 deficiency alone (Figure 4h, lane 3 vs 4). These data further positioned CLCC1 in lipoprotein biogenesis, prior to the transport step mediated by the SURF4 receptor. Consistently, quantitative measurements in liver weight, lipid levels, and plasma AST/ALT also supported the functional convergence between CLCC1 and the TMEM41B scramblase *in vivo* (Extended Data Figure 7b-e), as afore-unveiled by the molecular investigations in Figure 3.

## CLCC1 counters membrane dysfunction associated lipid stress

The preceding studies underscored severe pathological consequences of sustained leaflet imbalance at the ER, necessitating efficient equilibration via lipid scrambling across the ER bilayer. The identification of CLCC1 as a regulatory factor, however, implied the active mobilization of lipid scrambling mechanism to orchestrate a dynamic balance, thereby avoiding deleterious progression from a potentially widespread physiological event. In line with the notion, loss of hepatic CLCC1 was accompanied with substantial elevation of lipogenic enzymes, including Lipin1, FASN, ACC1. By contrast, ER chaperones for protein folding such as BIP and calnexin remained unaltered (Figure 5a). Interestingly, the lipid transfer enzyme complex MTP/PDI was also elevated, suggesting increased lipid transfer and therefore biased partition upon ER leaflet imbalance^33–35^. Indeed, MTP inhibition in CLCC1 deficient cells with pharmacological inhibitors^36,37^ redirected neutral lipids from geLDs marked by GFP-CB5 to small cLDs delineated by GFP-Plin2 (Figure 5b, quantified in c). Similar lipid redistribution was observed with MTP inhibition in TMEM41B deficient cells (Extended Data Figure 8a). The data thus indicated that phospholipid asymmetry between ER leaflets may be coupled with directional lipid transfer, furthering a physiological role of properly-regulated bilayer imbalance in lipid regulation.

**Figure 5.**
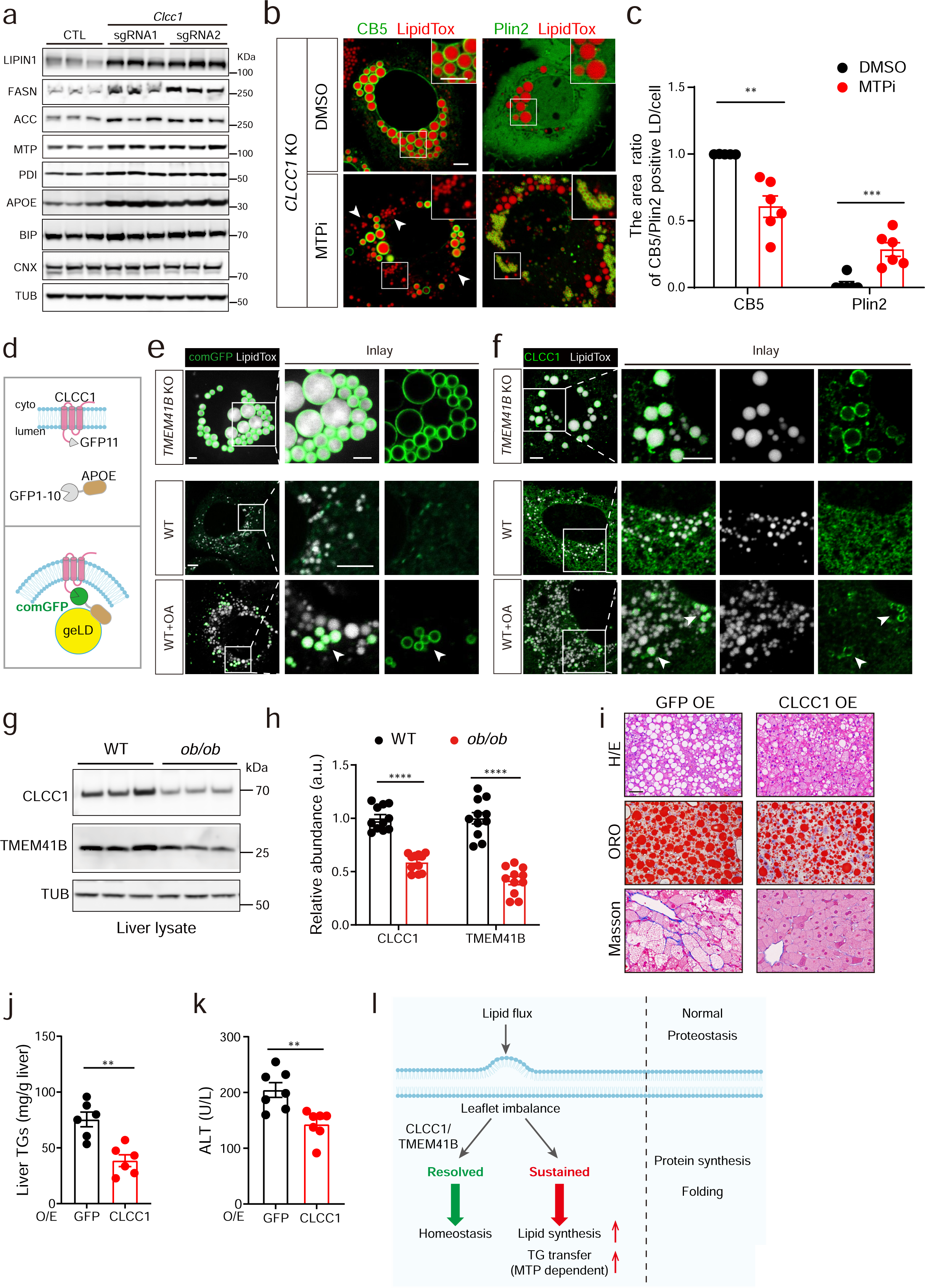
CLCC1 counters membrane dysfunction associated lipid stress. **a.** Loss of hepatic CLCC1 upregulates lipogenic and lipid transfer enzymes. IB analysis of liver samples from mice receiving indicated sgRNAs was conducted. **b.** MTP inhibition redirects neutral lipids from geLD to cLD. CRISPR/Cas9-mediated *CLCC1* KO Huh7 cells infected with AAVs expressing GFP-CB5 (left) or GFP-Plin2 (right) were treated with DMSO (upper) or MTP inhibitors (lower) for 24 h. Cells were stained with LipidTox prior to confocal microscopy. Scale bar, 5 μm. **c.** Quantification of cLD/total LD area from (b). Data are presented as mean ± SEM. **p<0.01, ***p<0.001 (two-tailed Student’s t test). n = 6 in each group. **d.** Schematic of CLCC1/ApoE complementation reporter. Upper, GFP11 was attached to the N-terminus of CLCC1, whereas GFP1-10 was fused to the C-terminus of ApoE. Lower, the engagement between CLCC1 and ApoE will complement the non-fluorescent GFP fragments, leading to fluorescence emission. **e.** GFP complementation in *TMEM41B* KO cells or WT cells treated with Oleic acid (OA). Huh7 cells with indicated genotypes or treatments were co-infected with AAVs expressing GFP11-CLCC1 or APOE-GFP1-10. At 16 h after infection, WT cells were treated with or without 200 μM OA for 8 h, followed by LipidTox staining and confocal microscopy analysis. Scale bar, 5 μm. **f.** CLCC1 re-localization upon OA treatment, inducing the enrichment of endogenous CLCC1 to encircle a subset of lipid droplets. *TMEM41B* KO Huh7 cells (upper) or WT Huh7 cells stimulated with control (middle) or 200 μM OA (lower) for 8 h, were co-stained with anti-CLCC1 antibody and LipidTox, prior to confocal microscopy. Scale bar, 5 μm. **g.** IB of hepatic CLCC1 and TMEM41B protein in *ob/ob* mice and lean control mice. **h.** Quantification of IB in (g). Data were analyzed from 3 independent experiments. Data are presented as mean ± SEM. ****p<0.0001 (two-tailed Student’s t test). **i.** Histological analysis of the livers from *ob/ob* mice receiving GFP or CLCC1. *Ob/ob* mice were injected AAVs containing TBG-GFP (control, n = 7) or TBG-CLCC1 (n = 7). Analysis was conducted at 4 weeks post AAVs administration. Scale bars: 50 μm. **j.** Quantification of liver TG in (i). Data are presented as mean ± SEM. **p<0.01 (two-tailed Student’s t test). **k.** Quantification of plasma ALT (i). Data are presented as mean ± SEM. **p<0.01 (two-tailed Student’s t test). **l.** Proposed model of leaflet imbalance and its equilibration constituting a homeostatic control mechanism intrinsic to ER function in lipid biogenesis and distribution. For **a-c, e-g,** representative results of at least 3 biological independent replicates are shown.

To further detect the general presence of leaflet imbalance and biased lipid partition, we designed a complementation-based reporter pair (Figure 5d). In brief, the non-fluorescent GFP fragments were fused to CLCC1 and the exchangeable apolipoprotein APOE (CLCC1-GFP11 and APOE-GFP1-10), respectively. Neither protein of the pair emitted fluorescence when expressed alone (Extended Data Figure 8b). However, when co-introduced into TMEM41B-deficient cells, the pair efficiently complemented fluorescent GFP, delineating the geLD surface (Figure 5e, upper panel). Interestingly, lipogenic treatment such as oleic acid (OA) also elevated the GFP signal complemented by the reporter pair in wild type cell (Figure 5e, middle and lower panels). Moreover, OA treatment relocated endogenous CLCC1 to encircle a small number of lipid droplets (Figure 5f, middle and lower panels), which were devoid of Plin2 but colocalized with GFP-CB5 and APOE-GFP (Extended Data Figure 8c-e). These data indicated potentially widespread leaflet imbalance at the ER during trans-bilayer lipid shuttle, furthering that lipid scrambling represent an actively regulated biological process to maintain a dynamic equilibrium.

To further investigate the general presence of ER leaflet imbalance and the metabolic defects resulted from sustained bilayer disequilibrium, we employed high fat diet-induced (HFD) or genetical predisposed obese mice (*ob/ob* mice). Interestingly, hepatic levels of CLCC1 protein were down-regulated by ∼50% in both HFD (Extended data Figure 9a&b) or *ob/ob* mice (Figure 5g, quantified in h). In line with the coincided loss of TMEM41B protein upon *Clcc1* inactivation observed in figure 2G, down-regulation of TMEM41B protein levels was also observed (Figure 5g&h), further implicating ER leaflet imbalance in the metabolic defects in these obese models. Interestingly, hepatic expression of CLCC1 alleviated lipid accumulation in *ob/ob* mice, as revealed by H/E, oil red staining, and hepatic lipid measurements (Figure 5i&j, Extended Data Figure 9c). Compared to GFP control, CLCC1 expression also ameliorated liver damages, as reflected by decreased liver enzymes leaked into the plasma (Figure 5k and Extended Figure 9d). Proteins levels of TMEM41B were also elevated by CLCC1 introduction, coincided with reduced lipogenic enzymes such as Lipin1 (Extended Data Figure 9e). In conclusion, our study established the poorly characterized ER membrane protein CLCC1 as an essential regulatory factor in scramblase-catalyzed bilayer equilibration at the ER, thereby actively counteracting widespread leaflet imbalance intrinsic to normal ER physiology in lipid biogenesis and distribution. Inadequate lipid scrambling, or sustained ER bilayer disequilibrium, triggers severe cellular and systemic pathologies despite normal proteostasis (Figure 5l). Considering the emergence of TMEM41B, CLCC1, and APOE in various diseases, spanning viral, neurodegenerative, and cardio-metabolic conditions^14,38–41^, future investigations may identify additional novel regulators to establish a common mechanism in lipid-based ER homeostatic control in both health and diseases.

## Supporting information

Extended Data Video 1, Three-dimensional rendering of tomogram corresponding to the region of WT hepatocytes in Figure 1b (left).

Extended Data Video 2, Three-dimensional rendering of tomogram corresponding to the region of Tmem41b KO hepatocytes in Figure 1b (right).

## Acknowledgments

The authors wish to thank Dr. Y. Jia for the anti-CLCC1 antibody and Drs. L. Chen and N. Gao for helpful discussions. The work is supported by National Science Foundation of China (NSFC) grants 92254308, 32125021, 92157107, 92357302 and National Key R&D Program grant no. 2021YFA0804802 and 2022YFA0806502.

## Note added in proof

During the preparation of this manuscript an additional, complementary characterization of CLCC1 in hepatic lipid flux was performed. Please refer to the corresponding preprint - Mathiowetz et al., “CLCC1 promotes hepatic neutral lipid flux and nuclear pore complex assembly.”.

## Methods

### Mouse models

All animal housing and use procedures were approved by the Institutional Animal Care and Use Committees of Peking University, an AAALAC accredited laboratory animal facility. All mice used in the experiments were bred on the C57BL/6J background. *Surf4 ^fl/lf^*mice were generated and maintained as previously described^24^. *Tmem41b ^fl/fl^* mice (RRID: IMSR_GPT: T026750) were generated by GemPharmatech (Nanjing, China) on the C57BL/6J background with loxP sites flanking exons 2 and 5, using CRISPR/Cas9 genome editing technology. Cre-dependent *spCas9* knockin (KI) mice were purchased from the Jackson Lab (RRID: IMSR_JAX: 026556)^26^. *Clcc1* LKO mice were obtained by intravenous injecting *spCas9* mice with AAV8 carring TBG-Cre and *Clcc1* targeting sgRNA. *Surf4^fl/fl^* mice and *Tmem41b^fl/fl^* mice were bred with with *spCas9* KI mice to generate *Surf4^fl/fl^*, *spCas9* mice and *Tmem41b^fl/fl^, spCas9* mice, respectively. Primers for the genotypes are listed in Extended Data Table 1. C57BL/6J mice were provided by Peking University at 6 weeks of age, and *ob/ob* mice (RRID: IMSR_GPT: T001461) were purchased from GemPharmatech (Nanjing, China) at 4 weeks of age. Mice were housed under standardized conditions, including a temperature of approximately 22°C, a 12 h light/dark cycle, and humidity of 40%-60%. Mice had free access to food and water unless otherwise stated. Male mice aged 6-16 weeks were used in all experiments. Mice were randomly assigned to different experimental groups. For fasting studies, all experiments were performed at 9 a.m. on the second day, with fasting starting at 5 p.m. on the first day.

### Cell culture

HEK293T cells (ATCC, CRL-3216), Huh7 cells (JCRB Cell Ban, JCRB0403), and HEK293F cells (Thermo Fisher Scientific, R79007) were obtained from ATCC, JCRB, and Thermo Fisher Scientific, respectively. HEK293F suspension cells were cultured in SMM 293T-I medium (Sino Biological, M293T1), supplemented with 0.5% penicillin/streptomycin (P/S) (Caisson, PSL01) and 1% fetal bovine serum (FBS) (VisTech, SE100-011) and maintained at 37°C in a 5% CO_2_ environment, with an optimal spinner speed of approximately 100 rpm to 130 rpm. The other cell lines were cultured in Dulbecco’s modified Eagle’s medium (DMEM) (HyClone, SH30022.01B) supplemented with 1% P/S and 10% FBS; under the same temperature and CO_2_ conditions. Transfections were performed with polyethyleneimine (PEI) (Polysciences, 23966-1) according to the manufacturer’s protocol.

### DNA vector construction

The sgRNAs were designed using the Benchling platform (https://benchling.com/) to optimize editing efficiency and minimize the unintended off-target effects. The sgRNA sequences are listed in the Extended Data Table 2. Oligonucleotides were cloned into either the pX602-AAV-Cre sgRNA backbone for CRISPR/Cas9-induced acute gene knockout in mouse liver or the pLentiCRISPR V2 (Addgene, 52961) & pLentiGuide Blast for genome editing in cell lines, following established protocols as described in previous research^42^. The pX602-AAV-Cre sgRNA construct was derived from the pX602 vector (Addgene, 61593) in our previous study^43^. The pLentiGuide Blast construct was derived from the pLentiGuide Puro vector (Addgene, 52963), in which blasticidin S deaminase was cloned between the BsiWI and MluI restriction sites to replace puromycin N-acetyltransferase.

pAAV-TBG-mCLCC1-StrepTagII-FLAG-StrepTagII was generated from pAAV-TBG-EGFP (Addgene, 105535) by replacing the GFP sequence with the murine *Clcc1* cDNA. For rescue experiments, the PAM region of m*Clcc1* cDNA was synonymously mutated as annotated in Extended Data Figure 3c.

For fluorescence imaging, mEGFP-Perilipin2, mEGFP-CB5 (residues 90 to 124 at the carboxyl-terminal end of cytochrome B5 type A, CYB5A), mEGFP-Sec61β, IgH signal peptide-mEGFP-KDEL, APOE-mEGFP, ApoE-GFP1-10, and GFP11-mCLCC1 (GFP11 was inserted into the N-terminus of mCLCC1, followed by the signal peptide) were cloned between KpnI and HindIII restriction sites to substitute for enhanced GFP (EGFP) in the pAAV-CAG-EGFP vector (Addgene, 51502).

For protein purification, CLCC1-FLAG, twin StrepTagII-TMEM41B and FLAG-TMEM41B were cloned into pKH3 (Addgene, 12555) between the EcoRI and XbaI restriction sites.

### Recombinant Adeno-Associated Virus (AAV) Production and Delivery

AAV packaging and purification were performed as previously described^43^. For the mouse experiment, AAV shuttle plasmids, Rep/Cap (2/8) plasmids, and helper plasmids were transfected into HEK293T cells using PEI. After 60 h post transfection, cells were harvested and virus was purified, quantified by both Coomassie Blue R250 staining and qPCR. AAV-2/8 was administered by tail vein injection. To achieve acute inactivation of the hepatic *Clcc1* gene in 6-week-old *spCas9* KI mice, each mouse was administered with pX602-AAV-Cre sgRNA at a viral genome copy number of 4×10^11^. For rescue experiments, each mouse was simultaneously injected with 4×10^11^ viral genome copies of pX602-AAV-Cre-sgRNA and 5×10^10^ viral genome copies of AAV-TBG-mCLCC1-StrepTagII-Flag-StrepTagII.

For AAV delivery into a cultured cell line, pRep/Cap (2/8) was replaced by pAAV-DJ for AAV-DJ preparation. AAJ-DJ with 1×10^10^ genome copies number was used per well in 6-well plate to delivery indicated gene into cultured cells. Subsequent experiments were conducted 24 h after AAJ-DJ infection, unless otherwise specified.

### Lentiviral packaging and knockout cell construction

HEK293T cells were used for lentivirus packaging. Briefly, the lentivirus shuttle plasmids, psPAX2 (Addgene, 12260), and pMD2.G (Addgene, 12259) plasmids were introduced into the cells using PEI according to the protocol provided by the manufacturer. Fouty-eight hours post transfection, the medium containing the lentivirus was collected and subsequently added to the WT Huh7 cells. Transducted cells were selected by antibiotic at 72 h after infection.

For control, *TMEM41B* KO and *CLCC1* KO Huh7 cell construction, pLentiCRISPR V2 lentivirus containing *Lacz* targeting sgRNA, human *TMEM41B* targeting sgRNA and human *CLCC1* targeting sgRNA (extended data table 2) were generated and infected into Huh7 cells, respectively. Two days after transfection, cells were selected with puromycin (Sigma, P8833). For TMEM41B & CLCC1 double deficient Huh7 cells, *TMEM41B* KO Huh7 cells were infected with pLentiGuide Blast lentivirus containing control or human *CLCC1* targeting sgRNA, and further selected with blasticidin (Wako, 022-18713).

### Plasma characterization and fast-protein liquid chromatography (FPLC) analysis

Blood samples were collected from the tail tips of mice fasted for 16 h using a heparinized capillary. Plasma was separated by centrifugation at 6000 rpm for 10 min at 4°C. Triglyceride, total cholesterol, ALT/GPT, and AST/GOT levels were determined using specific commercial kits (Sigma, TR0100; 000180, 000000010, 000000020 from Zhongsheng beikong, respectively) according to the manufacturer’s instructions. For FPLC analysis, pooled plasma samples from the same treatment group were fractionated using Superose 6 increase columns. The fractions were collected at a flow rate of 0.5 mL/min for subsequent measurements of cholesterol and triglyceride levels.

### VLDL secretion assay

Tyloxapol (Sigma, T8761) was administered to mice at a dose of 50 mg/kg body weight after a 16 h fast. Blood samples were collected at 1, 2, and 4 h after injection, and plasma triglyceride levels were analyzed as previously described.

### Histology

Tissue samples were collected and preserved in 4% paraformaldehyde (PFA) in PBS (Leagene, DF0135). Tissue embedding, sectioning, and hematoxylin and eosin (H&E) staining were performed by the Pathology Center of Peking University or Beijing ZKWB-Bio Biotechnology Co., Ltd. For oil red O staining, tissues were embedded in OCT compound and rapidly frozen. Cryosections of 8 μm thickness were cut and stained with Oil Red O according to the manufacturer’s instructions. Immunohistochemistry was performed on paraffin-embedded liver sections. Sections were deparaffinized and rehydrated, followed by antigen retrieval. To prevent nonspecific binding, sections were blocked with 10% goat serum for 1h and then incubated overnight at 4°C with primary antibodies diluted in blocking buffer. The sections were then exposed to secondary antibodies for 2 h at room temperature. Finally, they were visualized by DAB (3,3’-diaminobenzidine) staining.

### Quantification of hepatic triglycerides and cholesterol

Liver samples were quantified and homogenized in PBS. Lipids were extracted from the homogenates according to the established protocols of the modified Bligh-Dyer method. Briefly, the homogenates were vigorously mixed with a chloroform-methanol mixture (2:1). After centrifugation, the organic phase was carefully collected and concentrated using a rotary evaporator under vacuum. The lipid extract obtained was reconstituted in a solution of 15% Triton X-100 (Sigma, X100) in ddH_2_O. Quantification of triglycerides and cholesterol was performed as previously described.

### LD isolation and identification

The procedure of LD isolation was modified from a previously published nature protocol^44^. Before LD dissection, mice were fasted for 4 h. After anesthesia, the liver was perfused through the portal vein to remove blood. The liver was cut into approximately 1 mm^3^ pieces in Buffer A (25 mM Tricine pH 7.6, 250 mM sucrose) plus 0.2 mM PMSF and protease inhibitors (Roche, 4693132001) and homogenized 10 times on ice with a Dounce type glass-Teflon homogenizer. The lysate was centrifuged twice at 500g for 10 min at 4 °C to remove debris. The postnuclear supernatant (PNS, 10 mL) was then transferred to a SW40 Ti centrifuge tube, and 2 mL Buffer B (20LmM HEPES pHL7.4, 100 mM KCl and 2LmM MgCl_2_) was added to the PNS. The tubes were loaded into a Beckman SW40 rotor, and centrifuged at 300,000g for 2 h at 4°C. After the ultracentrifugation step, the white band of LD floating on the top of the solution was carefully collected into a new SW40 Ti centrifuge tube filled with 12 mL Buffer B, and centrifuged at 300,000g for 30 min, to remove contaminating organelles. This was repeated for three times to wash the isolated LD until no more pellets could be observed. LD proteins were precipitated with acetone at −80°C overnight, centrifuged at 15,000g for 10 min, washed with ice-cold acetone, dried with the lid open, and then solubilized in RIPA buffer (50 mM Tris-HCl pHL7.4, 150LmM NaCl, 1% Nonidet P-40, 1% sodium deoxycholate, 0.1% SDS, 1 mM EDTA, 10% glycerol with protease inhibitor) supplemented with 2% SDS. Protein concentrations were measured using the BCA Protein Assay Kit (Thermo-Pierce, 23227). Silver staining of LD proteins was performed with a commercial kit according to the manufacturer’s protocol (Thermo Scientific, 24600). Proteomics analysis was performed according to a previously published procedure (Wang et al., 2021). The proteins with more than 1 unique peptides were further analyzed. The criterion for identifying differentially expressed proteins were a fold change greater than 10. Enrichment analysis of upregulated or downregulated different expressed proteins was performed separately using Cluster Profiler R package (version 4.10.0). Venn analysis was performed using the VennDiagram R package (version 1.7.3). The significant GO terms of the cellular component were visualized using the ggplot2 R package (version 3.4.4).

### Immunoblotting

Liver lysates were prepared using RIPA buffer supplemented with protease inhibitors. Protein concentrations were measured using the BCA Protein Assay Kit. 1× SDS/PAGE sample buffer was added. Equal amounts of protein were loaded onto 3% to 15% Tris-acetate SDS/PAGE gels and transferred overnight at 4°C to nitrocellulose membranes (Cytiva, 10600006). The membranes were then blocked with 5% milk in TBSTL(20 mM Tris pH 7.4, 150 mM NaCl and 0.1% Tween 20) for 1 h at room temperature. Primary antibodies were diluted in TBST containing 5% milk and 0.02% Na_3_N, incubated with membranes overnight at 4°C and washed 3 times with TBST at room temperature, for 15 min each time. HRP-conjugated goat secondary antibodies were diluted in TBST containing 5% milk and incubated for 1 h at room temperature, followed by 3 washes. Membranes were visualized after exposure to ECL substrate (Thermo Fisher, 34578).

The following primary antibodies were used for immunoblotting: rabbit polyclonal anti-Perilipin2 (Cell Signaling Technology, 45535, 1:1,000); rabbit polyclonal anti-HSD17B13 (ABclonal, A65256, 1:1,000); rabbit polyclonal anti-ATGL (Cell Signaling Technology, 2138, 1:1,000); rabbit polyclonal anti-SigmaR1 (Proteintech, 15168-1-AP, 1:1,000); rabbit polyclonal anti-Calnexin (Proteintech, 10427-2-AP, 1:1,000); rabbit polyclonal anti-ApoB (Proteintech, 20578-1-AP, 1:1,000); rabbit polyclonal anti-ApoE,(Fitzgerald, 10R-10633, 1:1,000); rabbit polyclonal anti-ApoA1 (Fitzgerald, 70R-15769, 1:1,000); rabbit polyclonal anti-LIPIN1 (Proteintech, 27026-1-AP, 1:1,000); rabbit polyclonal anti-FASN (Proteintech, 10624-2-AP, 1:1,000); rabbit polyclonal anti-ACC1 (Proteintech, 21923-1-AP, 1:1,000); mouse monoclonal anti-MTP (Santa Cruz Biotechnology, sc-135994, 1:1,000); mouse monoclonal anti-PDI (BD transduction LaboratoriesTM, 610946, 1:1,000); rabbit polyclonal anti-BIP (Proteintech, 11587-1-AP, 1:1,000); rabbit polyclonal anti-Tublin (Proteintech, 11224-1-AP, 1:1,000). Rabbit polyclonal antibody against the N-terminal epitope of human TMEM41B (residues 1-109) were produced by the Proteintech Group (www.ptglab.com), the rabbit anti-serum was collected and purified by NHS beads conjugated with His-tagged human TMEM41B (redisues 1-109), followed by washing with PBS containing 0.15% Triton X-100, eluted with 50 mM glycine (pH 2.5) and neutralized with Tris-HCl pH 8.0. Rabbit polyclonal antibody against the C-terminal epitope of mouse CLCC1 (residues 355-539) was a gift from Dr. Yichang Jia.

The following secondary antibodies were used: goat anti-mouse IgG (H+L) secondary antibody (Thermo Fisher, 31430, 1:10,000) and goat anti-rabbit IgG (H+L) secondary antibody (Thermo Fisher, 31460, 1:10,000).

### Blue Native PAGE

Blue native PAGE analysis of CLCC1followed from a protocol described in 2006^45^. In brief, livers from the indicated genotype mice were disrupted in Buffer C (25 mM Tris pH 7.4, 250 mM sucrose, 1 mM EDTA and protease inhibitors) and homogenized 10 times on ice using a Dounce type glass-Teflon homogenizer. The lysate was centrifuged at 1000g for 10 min at 4 °C to remove nuclei and unbroken cells. The supernatants were further centrifuged at 100,000g for 30 min to isolate membrane fractions. Membranes from indicated mice and geLD from *Tmem41b* LKO mice were incubated with Buffer D (25 mM Tris pH 7.4, 50 mM NaCl, 1% n-dodecyl-ß-D-maltoside (DDM, Qisong biological, QS81007015) and protease) for 2 h at 4 °C, followed by centrifugation at 100,000g for 30 min. The concentrations of the supernatants were measured using the BCA Protein Assay Kit. 5% Coomassie Brilliant Blue G-250 dye was added at a final dye to detergent ratio of 1/8 (g/g). Equal amounts of protein were loaded on 4-16% blue native PAGE gel, transferred overnight at 4 °C to PVDF membranes (Cytiva, 10600023), followed by IB analysis.

### Immunoprecipitation (IP)

For immunoprecipitation of endogenous TMEM41B, cells were washed twice with ice-cold PBS and incubated with lysis buffer (50 mM Tris-HCl pHL7.4, 150 mM NaCl, 1% Nonidet P-40, 1 mM EDTA, 10% glycerol with protease inhibitor) for 20 min on ice. The lysate was centrifuged at 13000 rpm for 15 min at 4 °C. Supernatants were collected and protein concentrations were measured using the BCA Protein Assay Kit. Supernatants were incubated with anti-TMEM41B antibody with end-over-end rotation for 2 h at 4 °C, followed by incubation with rProtein A beads 4FF (Smart-life sciences, SA015025), continuously rotated for 4 h at 4 °C. The beads were washed five times with wash buffer (50 mM Tris-HCl pHL7.4, 150 mM NaCl, 0.1% Nonidet P-40, and 10% glycerol). For immunoblot analysis, proteins were further eluted from the beads with elution buffer (50 mM Tris-HCl pHL7.4, 150 mM NaCl, 0.1% Nonidet P-40, 1 mM EDTA, 10% glycerol, protease inhibitor with 0.4 mg/mL TMEM41B (residues 1-51) peptides). After gel separation, proteins were transferred to NC membrane for IB.

### Protein expression and purification

HEK293F cells were transfected with CLCC1-FLAG, FLAG-TMEM41B or CLCC1-FLAG & twin StrepTagII-TMEM41B plasmids. Cells were grown for 48 h, and harvested by centrifugation at 1,000g for 10 min at 4 °C, then washed with ice-cold TBS buffer. Cell pellets were resuspended in Buffer E (50 mM Tris-HCl pHL7.4, 150 mM NaCl, 1% DDM, 1 mM EDTA, 10% glycerol with protease inhibitor). After 2 h incubation, the cell lysate was spun at 100,000g for 30 min, and the supernatant was incubated with prewashed anti-DYDDDDK affinity beads (Smart-lifesciences, SA042500) for 2 h at 4 °C, the beads were washed with Buffer E (50 mM Tris-HCl pHL7.4, 150LmM NaCl, 0.025% DDM and 10% glycerol). The protein was eluted with Buffer E containing 0.4 mg/mL Flag peptides. For tandem purification of the CLCC1-TMEM41B complex, the flag peptide-eluted products were further incubated with streptactin beads 4FF (Smart-lifesciences, SA053500) for 2 h at 4 °C, the slurry was washed, and eluted with Buffer E containing 10 mM desthiobiotin (Sigma, 71610-M). Proteins were loaded onto Amicon 0.5 mL concentrators (10 KDa cutoffs, SEP, UFC501008), concentrated, buffer replaced with Buffer E, and quantified by Coomassie Blue R250 staining using BSA standards.

### Immunofluorescence and confocal microscopy

Cells were plated in 35mm glass bottom dishes (Cellvis, 150680) for subsequent treatment and staining.

For immunofluorescence, cells were fixed with 4% PFA in PBS for 10 min at room temperature. After fixation, cells were washed three times with PBS, then permeabilized with Immunostaining Permeabilization Buffer with Saponin (Beyotime, P0095) for 20 min and washed three times with PBS. After blocking with 2% BSA in PBS for 1 h, the cells were incubated with primary antibodies in blocking solution overnight at 4 °C. The cells were then washed three times with PBS. The cells were then incubated with fluorescence-labeled secondary antibodies for 1 h at room temperature and washed three times with PBS. For staining, lipid droplets were visualized with HCS LipidTox^TM^ Red (Thermo Fisher, H34476, 1:1,000) or HCS LipidTox^TM^ Deep Red (Thermo Fisher, H34477, 1:1,000) for 20 min, and immersed in PBS or complete cell culture medium for imaging.

The following primary antibodies were used for immunofluorescence: mouse monoclonal anti-FLAG (M2) (Sigma, F1804, 1:500); rabbit polyclonal antibody against the C-terminal epitope of human CLCC1 (residues 355-539) (1:500), a gift from Dr. Yichang Jia; rabbit polyclonal anti-TMEM41B (1:20), purified from rabbit serum against TMEM41B (residues 1-109) using the N-terminal 1-20 amino acids peptide; Goat anti-Rabbit IgG (H+L) Cross-Adsorbed Secondary Antibody, Alexa Fluor^TM^ 488 (Thermo Fisher, A-11008, 1:1,000); Goat anti-Rabbit IgG (H+L) Cross-Adsorbed Secondary Antibody, Alexa Fluor^TM^ 568 (Thermo Fisher, A-11011, 1:1,000); Goat anti-Rabbit IgG (H+L) Cross-Adsorbed Secondary Antibody, Alexa Fluor^TM^ 568 (Thermo Fisher, A-11031, 1:1,000).

Images were acquired on a ZEISS 900 confocal microscope with Airyscan2 (Carl Zeiss) and processed with ImageJ (NIH).

### Image analysis

To analyze TMEM41B signal enriched around LD, images were imported into ImageJ (NIH). LD ROIs were detected by “Threshold-Analyze particle” and the mean intensity of TMEM41B at LD ROIs was measured. Then, the mean intensity of TMEM41B at 5 cytosol ROIs was randomly measured. The ratio of mean intensity of LD ROIs to cytosol ROIs was defined as TMEM41B enrichment around LD.

To quantify percentages of LDs with Plin2 or CB5 signals, images were imported into ImageJ (NIH). LDs with Plin2 or CB5 were detected and collected by “Threshold-Analyze particles (include holes)” at correspond channel. And total LDs were detected “Threshold-Analyze particles” at LD’s channel. Then, LD number or Area with the Plin2 or CB5 signals divided total LD number or area are the percentages of LDs with Plin2 or CB5 signals.

### Association analyses of the SNPs with plasma lipid level

GWAS summary statistics of the *CLCC1* gene and plasma lipid levels were obtained from the GLGC dataset^25^ and performed using LocusZoom (http://locuszoom.org/). The allele frequency in humans of rs149700491 SNP was available from 1000 Genome Phase 3.

### *In vitro* lipid scrambling assay and NBD-Glucose leakiness assay

POPC (Sigma, 42773), POPG (Sigma, 840457P) and NBD-PC (Sigma, 810133P-1MG) were solubilized in chloroform, at a molecular ratio 180:19:1, the lipids were dried under an argon or nitrogen stream, and the flask was transfected to a vacuum desiccator at room temperature for at least 3 h. The dried lipid film was reconstituted in Buffer F (50 mM HEPES pH 7.4, 200 mM NaCl,) at a concentration of 5.25 mM lipid. This lipid solution was incubated in a water bath at 37°C for 10 minutes, subjected to ten freeze-thaw cycles, and extruded 30 times through a 400 nm polycarbonate filter (Avanti, Cat# 610020). Liposomes were destabilized with 7mM DDM at room temperature for 2 h, then detergent-solubilized protein preparations (TMEM41B, CLCC1, CLCC1-TMEM41B complex) and detergent buffer control were added at the indicated concentration, and incubated for 1 h. Bio-beads (BIO-RAD, 1528920) were used to remove detergent according to the manufacturer’s protocol. Samples were dialyzed in buffer F with a magnetic rotor at 4°C for 48 h.

The scramblase assay was performed in a cuvette equipped with a magnetic stirrer at room temperature, with 2 mL reaction volume (∼200 nM lipid concentration). Fluorescence signals were monitored (excitation at 460 nm, emission at 538 nm) using the Synergy H1 Hybrid Multi-Mode Reader (BioTek) at 1-second intervals. To assess lipid scrambling, NBD fluorescence was monitored over time after the addition of dithionite (Sigma, 71699) to a final concentration of 10 mM. Subsequently, an additional Triton X-100 (0.5%) was then added to disrupt the liposomes, allowing for a thorough reduction of all NBD fluorescence.

The NBD-glucose leakage assay was performed in a manner similar to the lipid scramblase assay described above, with the significant difference that NBD-PC was omitted from the liposome composition. Instead, NBD-glucose (Abcam, ab146200, 20 μM) was added to the buffer during the destabilization step. During the dialysis process, the NBD-glucose in the liposomes is preserved.

### Click labeling with alkyne-choline

CRISPR/Cas9-mediated control (CTL), *TMEM41B* KO, *CLCC1* KO, and *TMEM41B*/*CLCC1* double KO Huh7 cells were seeded on glass coverslips in 6-well plates at 25% density, and cultured with 100 μM alkyne-choline (CONFLUORE, BCP-44) for 18 h. Cells were fixed with 4% PFA in PBS for 10 min at room temperature and washed three times with PBS. The cells were then permeabilized with 25 μg/mL digitonin for 15 min and washed three times with PBS. Cells were incubated with 10 μM 5-TAMRA azide (Confluore, BDR-22), BTTAA (Nonidet P-40, BDJ-4)-CuSO4 complex (50 μM CuSO4, BTTAA/CuSO4 6:1, mol/mol) and 2.5LmM sodium ascorbate (Aladdin, S105024) in PBS at room temperature for 1 h, and washed three times with PBS. After the click reaction, coverslips were mounted on glass slides and examined under a ZEISS 900 confocal microscope with Airyscan2 (Carl Zeiss). Fluorescence intensity was quantified using ImageJ Fiji (NIH) and analyzed using Prism (GraphPad)

### EM samples preparation and images

Mice were euthanized and perfused systematically with 0.1 M sodium phosphate buffer (PB, pH 7.4), followed by a pre-fixation solution containing 2.5% glutaraldehyde and 0.8% paraformaldehyde. The liver tissues were then surgically excised, fixed in the pre-fixation solution for 2 h at room temperature, and further dissected into smaller sections (0.2 × 0.3 × 0.5 mm^3^). These samples underwent an additional overnight fixation at 4°C in the same pre-fixing solution. After rinsing in PB, the tissues were immersed in 0.1 M imidazole in 0.1 M PB for 30 min and then postfixed in 2% osmium tetroxide in 0.1 M PB. After thorough rinsing with ultrapure water, the tissues were stained with 1% uranyl acetate at 4°C overnight. The samples were dehydrated through a gradient acetone series, then embedded in epoxy resin, and polymerized at 60°C for 24h. The prepared samples were sectioned at approximately 60 nm thickness using a Leica EM UC7, placed on copper grids, and imaged using an FEI Tecnai G2 20 Twin electron microscope equipped with a CMOS camera (EMSIS, Germany). And the examination of liver tissue structure was carried out using a double-blind method.

### Cryo-ET Sample Preparation

The liver samples were vitrified using a previous established protocol^18^. In brief, following cervical dislocation, the abdomen of each mouse was dissected. A small piece of liver tissue was meticulously excised using a scalpel and placed onto an EM grid (Beijing XXBR: Cu microgrid, T10012, 200 mesh). The grid was coated with an additional carbon layer (∼10 nm thick) and freshly glow discharged using a Model 950 Advanced Plasma System (Gatan, Pleasanton, CA, USA) before usage. The grid was previously placed into a 6 mm aluminum carrier (200 nm in depth). After filling the carrier with 2-methylpentane (Sigma, M65807), a 6 mm sapphire disc was immediately placed on top of the carrier. The assembly of the sandwich carrier was promptly inserted into the high-pressure cryostat (Leica EM ICE) for vitrification. The vitrified sandwich assembly was then transferred to a liquid ethane and propane mixture (ethane:propane=36.9%:63.1%) at −170°C, allowing 2-methylpentane to dissolve. The resulting grid was then transferred to liquid nitrogen for storage until milling.

A cryo-FIB/SEM (Aquilos 2, Thermo Fisher Scientific) was employed to prepare lamella, using an adapted serial lift-out method^46^. Briefly, the grid was assembled in a ThermoFisher FIB Autogrid, before being loaded into the FIB chamber via the sample transfer rod. To minimize the ion beam damage and enhance the electrical conductivity, the sample was coated with a layer of Pt by GIS for 2 min, followed by sputtering at 30 mA for 15 s. Next, the target area of the sample was bombarded with the ion beam to isolate the sample chunk. This chunk was subsequently attached to the cryo-needle and transferred to the receiving EM grid (Beijing XXBR: Cu rectangular grid, G100/400, 100/400 mesh), where it was serially sectioned to several lamellae. These lamellae were securely attached to the grid in turn by redeposition method. Finally, each lamella was finely milled to a target thickness of approximal 200 nm, which was then used for subsequent tomographic data acquisition.

### Cryo-ET Data Acquisition

The lamellae from wild-type mice were loaded into a 300 kV cryo-transmission electron microscope Titan Krios G3 (ThermoFisher Scientific) equipped with a K3 camera and a Gatan energy filter. Automatic tomographic tilt series acquisition was performed using SerialEM software^47^. Images were acquired at a magnification of 64000× (pixel size was 1.37 Å) in MRC format using super-resolution mode, resulting in 10 frames per image. The defocus was set from −4.0 to −6.0 μm. The acquisition was performed from −50° to +50° (with respect to the pre-tilt angle) in 2° increments, utilizing the dose symmetric scheme^48^. The total electron dose was limited to 120 e/Å^2^ per tilt series.

For the lamellae from *Tmem41b* LKO mice, they were loaded to a 300 kV cryo-transmission electron microscope Titan Krios G4 (ThermoFisher Scientific) with a Falcon 4i camera and a ThermoFisher energy filter. Automatic tomographic tilt series acquisition was performed using Tomography 5.12.0 (ThermoFisher Scientific). Images were acquired at a magnification of 42000× (pixel size was 3.00 Å) in EER format. The defocus was set from −2.5 to −5.0 μm. The acquisition was performed from −50° to +50° (with respect to the pre-tilt angle) in 2° increments, utilizing the dose symmetric scheme^4^. The total electron dose per tilt series was limited to 120 e/Å^2^.

### Tomogram Reconstruction and Membrane Segmentation

TOMOMAN and TOM toolbox^49^ were used as general tools in image processing. Initially, all frames of each tilt were motion corrected using MotionCor2 software^50^. Subsequently, each tilt series was aligned using patch-tracking method in IMOD software^51^, and then reconstructed using back projection method to obtain a tomogram. For visualization, all tomograms were rescaled before further processing. Specifically, the wild-type tomograms were binned by a factor of 8, while tomograms from *Tmem41b* KO sample used a binning factor of 4 or 6 depending on the desired resolution. Then a deconvolution filter was applied to all tomograms to further improve the contrast. Membrane segmentation was firstly performed using MemBrain-Seg^52^, and then manually polished with Amira 3D 2022.2 (ThermoFisher Scientific). Lipid droplets were manually segmented with Amira. ChimeraX 1.7.1^53^ was used for the final rendering.

### CLCC1 structure prediction

The amino acid sequence 91-360 of human CLCC1 was used to predict the monomer and hexamer structure using AlphaFold3^30^.

### Quantification and statistical analysis

All experimental results are presented as mean ± SEM unless otherwise noted in the figure legends. Sample sizes were not predetermined by statistical methods. Statistical analysis was performed with GraphPad Prism 9. Statistical significance was determined using Student’s t-test or one-way ANOVA with Tukey’s post hoc test, as indicated in the figure legends. Results were considered significant if P < 0.05. Statistical significance levels are indicated in the figure legends. Experimental results shown are representative of at least 3 independent experiments. Mouse experiments were randomized. Imaging and histology were evaluated in a blinded fashion.

---

**Extended Data Figure 1.**
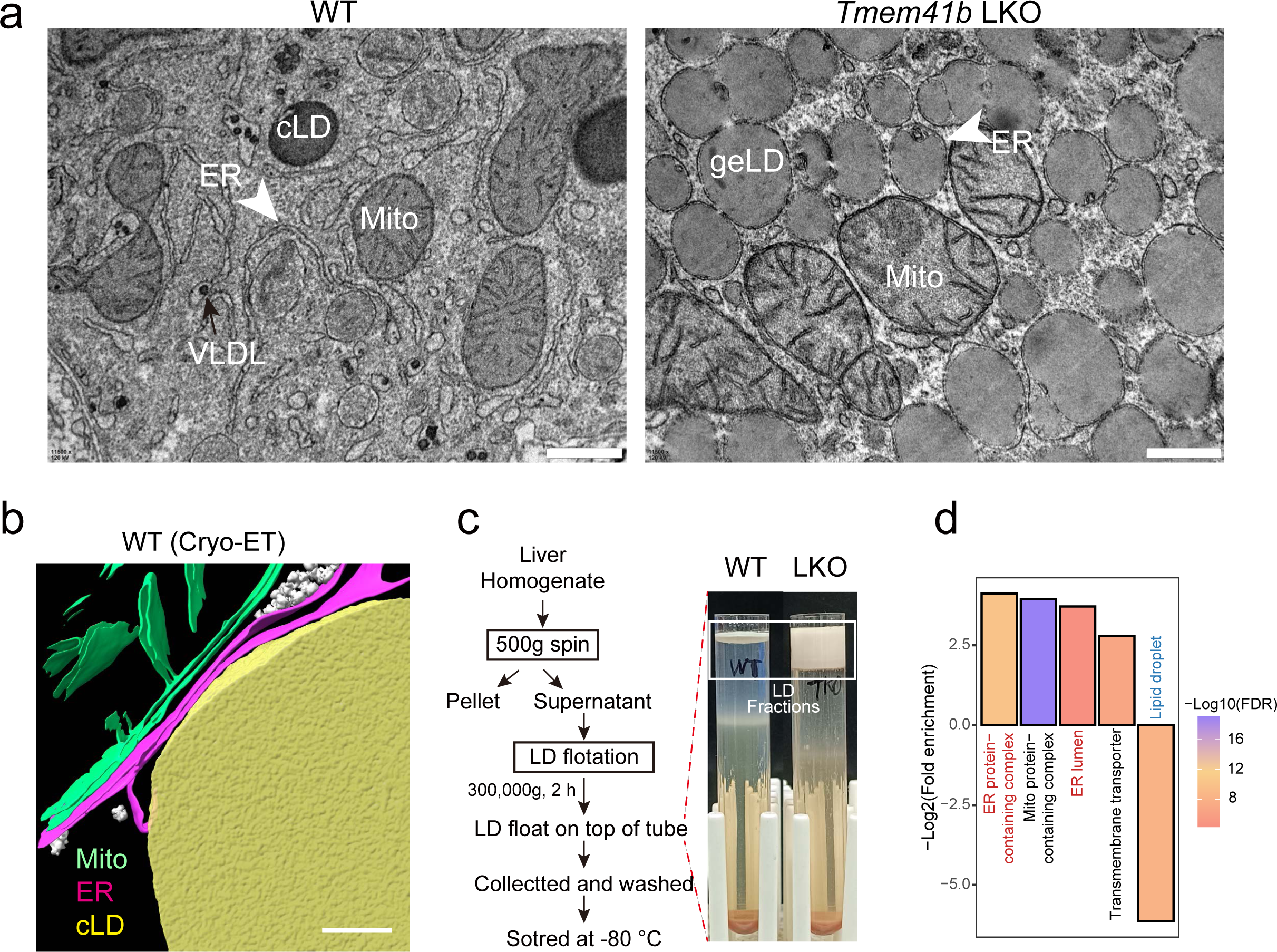
TMEM41B deficiency results in gaint ER-enclosed lipid droplets (geLD) a. TEM images of the livers from WT (left) or *Tmem41b* LKO (right) mice. Arrowheads indicate the ER. Scale bars, 500 nm. n = 5 mice for each genotype. b. Three-dimensional rendering of tomogram corresponding to the region of WT hepatocytes in Figure1b. Yellow: cLD, green: mitochondria, and magenta: the ER. Scale bars, 100 nm. c. Workflow of LD isolation from mouse liver (left). Images of cLD and geLD after density gradient fractionation (right). d. Gene ontology analysis of differential proteins in geLD compared to cLD. For c and d, representative results of at least 3 biological independent replicates are shown.

**Extended Data Figure 2.**
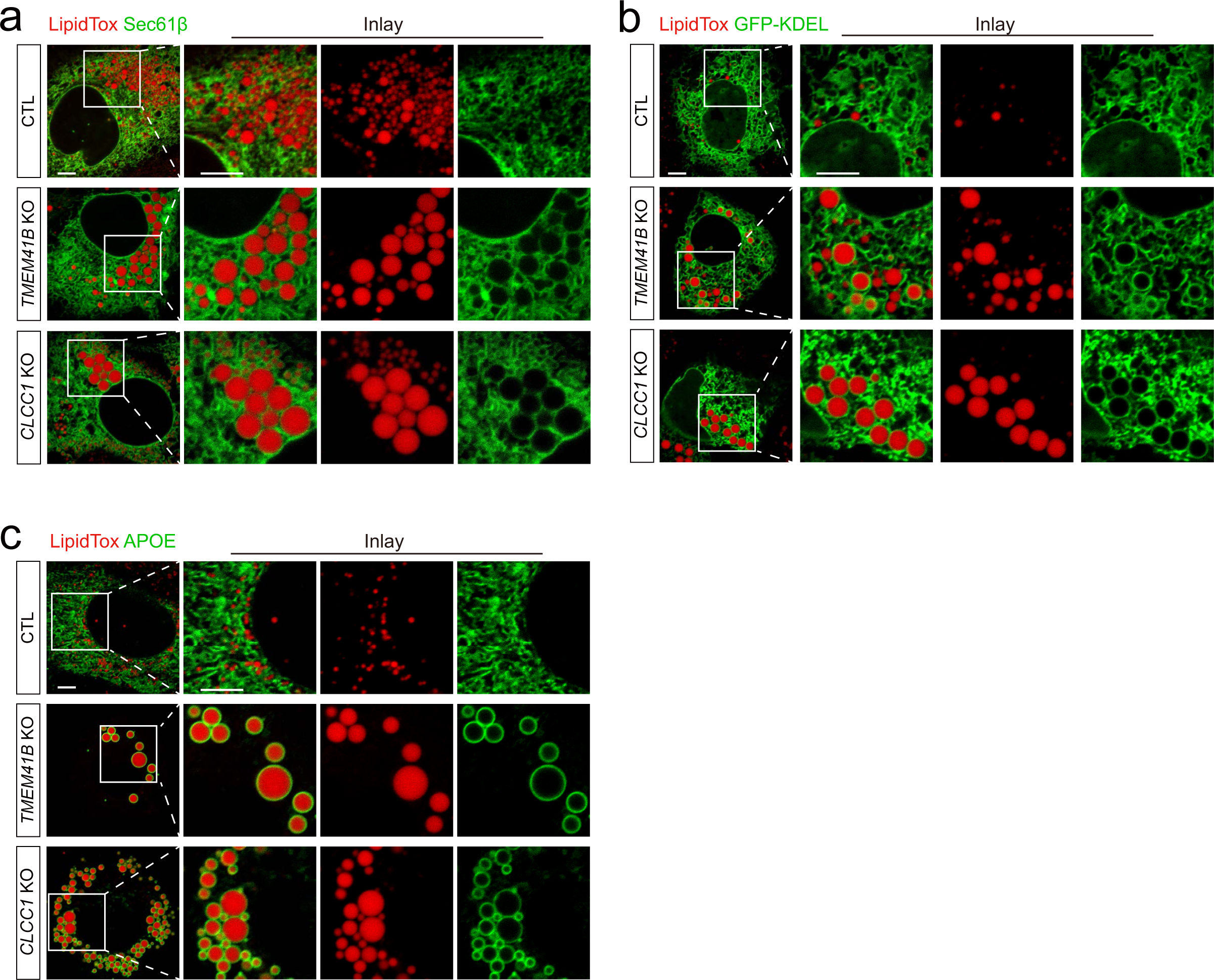
The geLDs are encircled by smooth ER bilayers in TMEM41B or CLCC1 deficient cells. a. Confocal microscopy of CRISPR/Cas9-mediated control (upper), *TMEM41B* KO (middle) and *CLCC1* KO (lower) Huh7 cells infected with AAVs expressing GFP-Sec61β. Red, LipidTOX. Scale bar, 5 μm. b. The same cells in (a) expressing GFP-KDEL. Red, LipidTOX. Scale bar, 5 μm. c. The same cells in (a) expressing ApoE-GFP. Red, LipidTOX. Scale bar, 5 μm. For a-c, representative results of at least 3 biological independent replicates are shown.

**Extended Data Figure 3.**
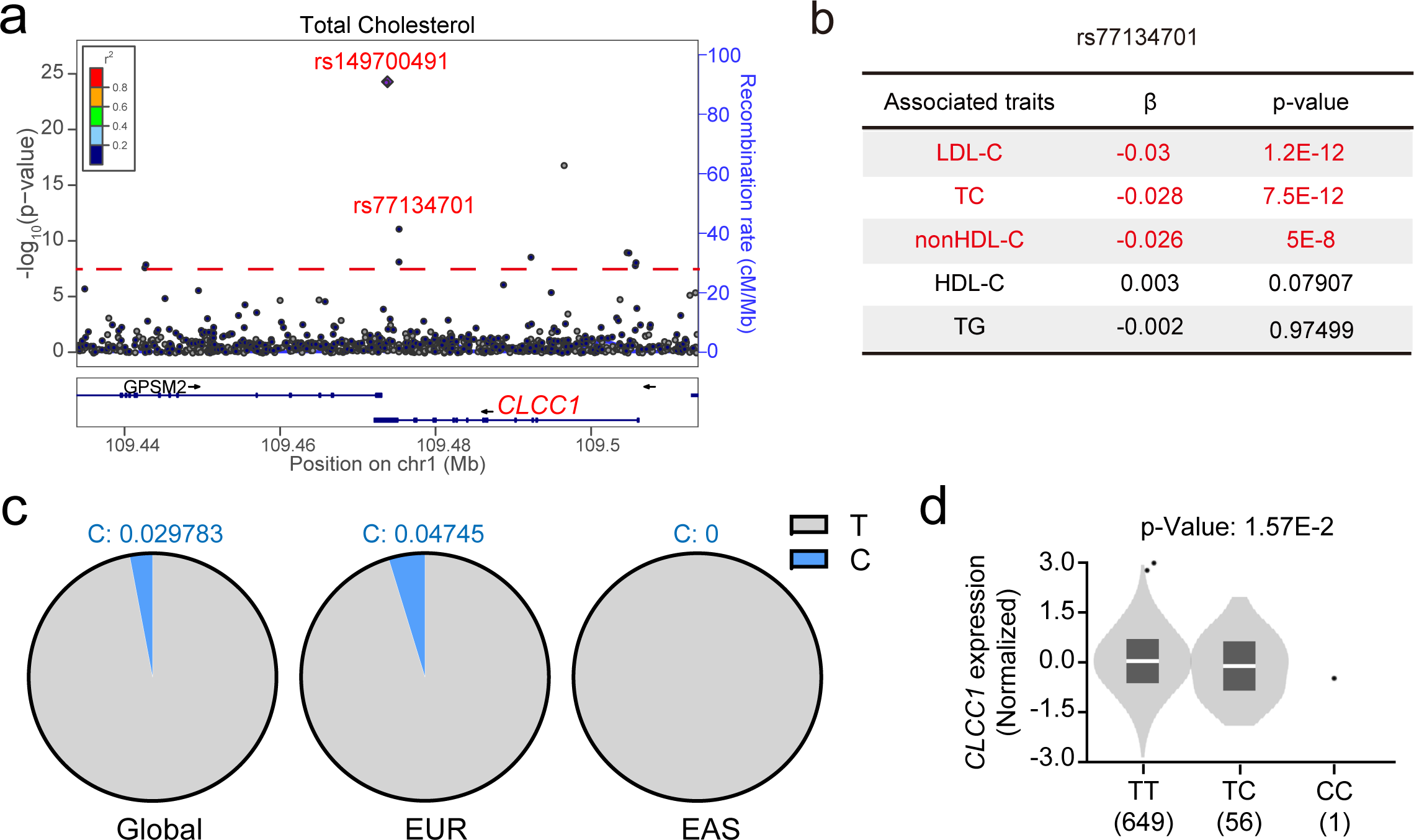
Genetic variants in the human *CLCC1* gene are associated with plasma lipids in populations. a. Regional plot of *CLCC1* associated with plasma LDL-cholesterol levels in humans. b. Summary of the GLGC genome-wide association data between the SNP rs77134701 in *CLCC1* and plasma lipids in humans. c. Minor allele frequency (MAF) difference at SNP rs77134701in European (EUR) and East Asian (EAS) descents from gnomAD-Genomes (Global). d. Minor allele of rs77134701 is associated with decreased *CLCC1* expression in human skeletal mus-cles. P-Value, 1.57E-2.

**Extended Data Figure 4.**
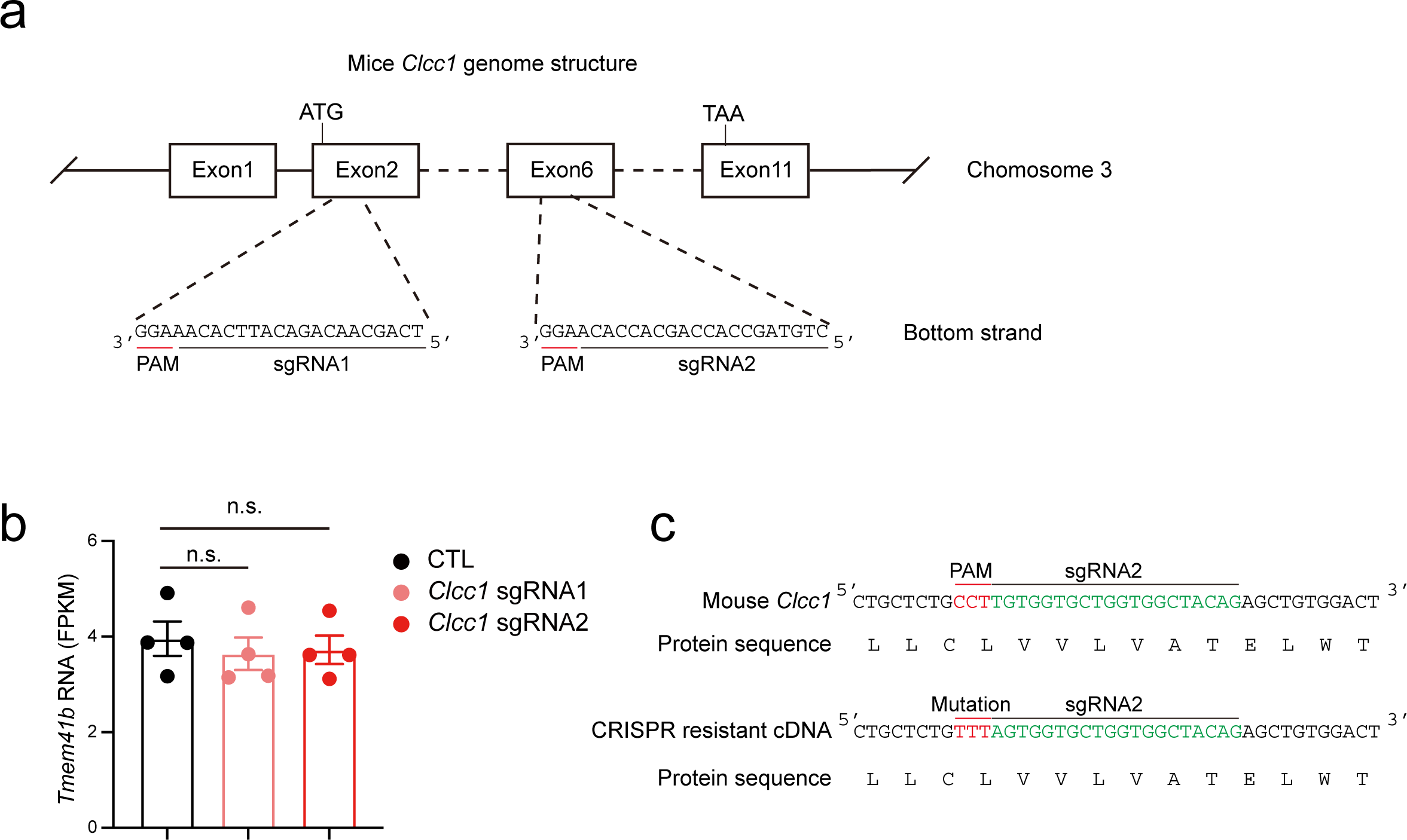
CRISPR/Cas9-mediated hepatic *Clcc1* knockout in mice. a. Schematic of *Clcc1* sgRNA1 and sgRNA2 targeting sites in the gene. b. The FKPM values of *Tmem41b* mRNA from RNA sequencing of *Lacz* sgRNA control (CTL), *Clcc1* sgRNA1 and *Clcc1* sgRNA2 mouse liver. n.s., not significant. c. Design of CRISPR-resistant *Clcc1* cDNA for the rescue experiments.

**Extended Data Figure 5.**
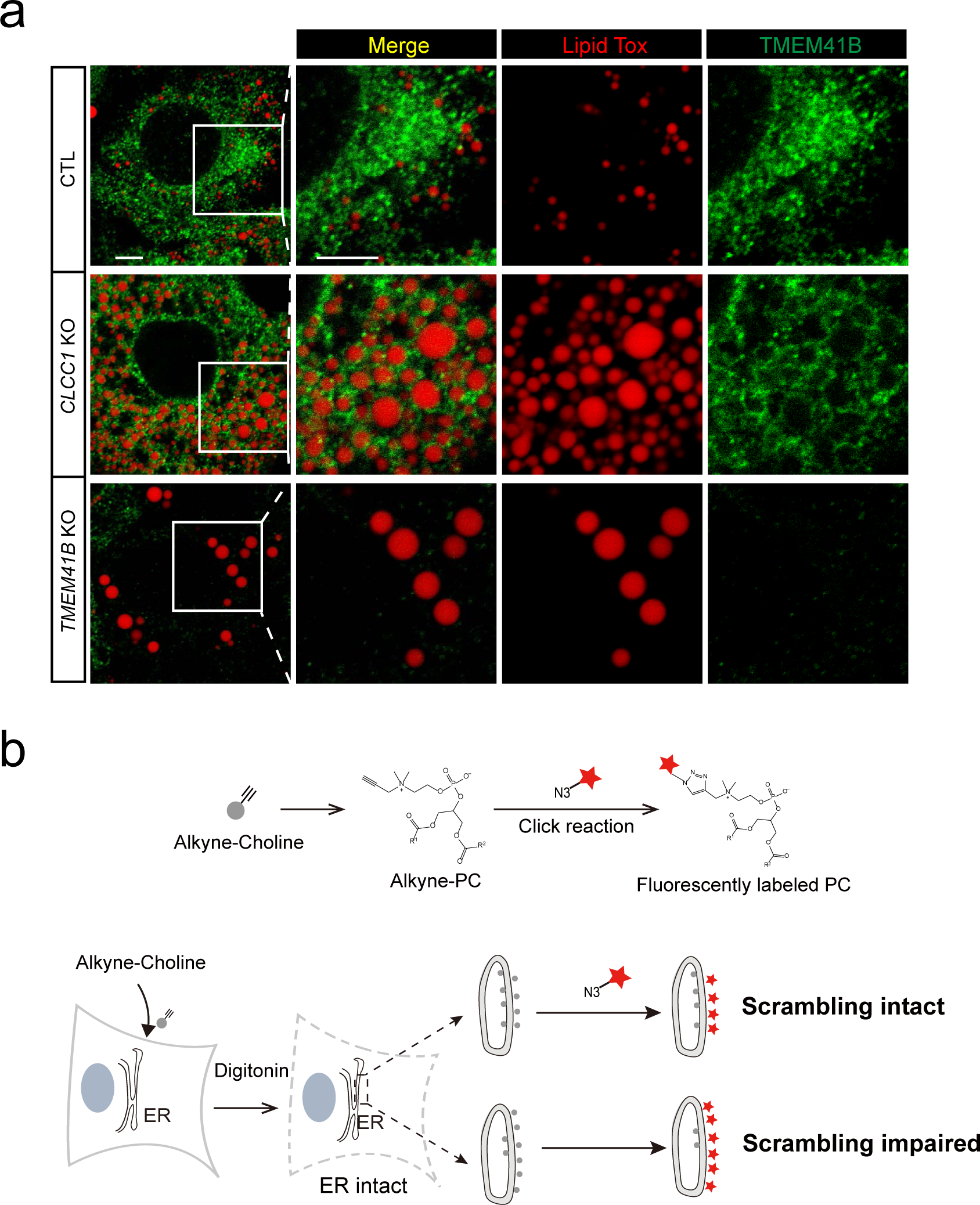
CLCC1 recognizes curved ER leaflets and regulates TMEM41B scram-blase activity. a. Little enrichment of endogenous TMEM41B around geLD in CLCC1 deficient cells. WT (upper), *CLCC1* KO (middle) and *TMEM41B* KO Huh7 cells were co-stained with anti-TMEM41B antibody and LipidTox, prior to confocal microscopy. Scale bar, 5 μm. Representative results of at least 3 biological independent replicates are shown. b. Schematic diagram of ER lipid scrambling assay in cells. Upper: a workflow of metabolic labeling and click-chemistry to detect newly-synthesized phosphatidylcholine (PC) in cells. Lower: Cells are metabolically labeled with alkyne-choline, allowing the synthesis of alkyne-PC at the cytosolic leaflet of the ER. In scrambling intact cells, alkyne-PC would be shuttled across the ER bilayer to the inner leaflet. Cell membranes are selectively permeabilized with Digitonin, without disrupting the ER bilayer. This procedure restricts ‘click-chemistry’ reagents to only label the alkyne-PC on the outer leaflet, but not the inner leaflet. When scrambling activity is impaired in cells, an accumulation of alkyne-PC occurs on the outer leaflet, resulting in enhanced fluorescent signal detection.

**Extended Data Figure 6.**
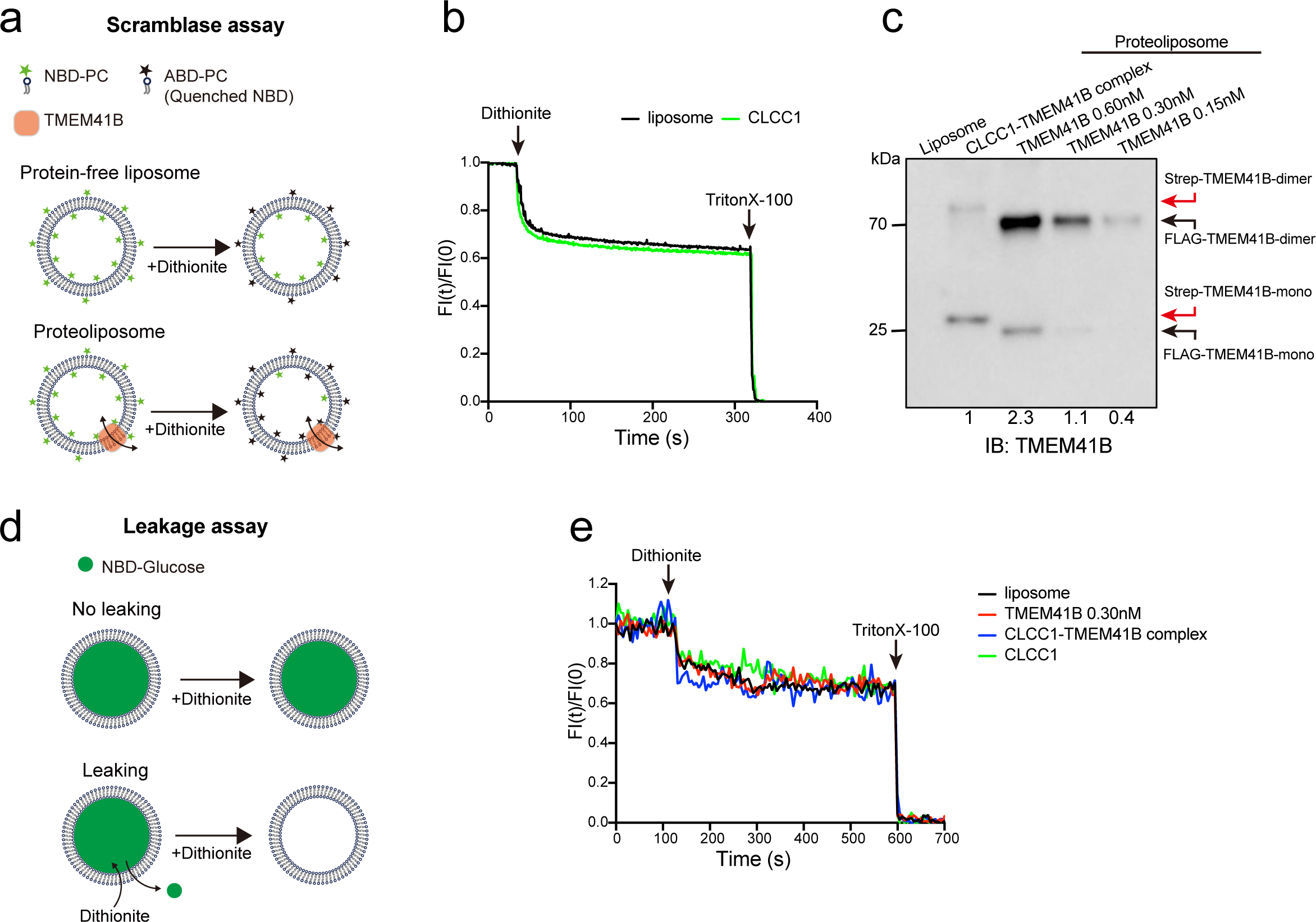
Scramblase assay and leakage assay *in vitro*. a. Schematic diagram of liposome-based lipid scramblase assay *in vitro*. NBD-PC is symmetrically distributed across both leaflets of liposomes. Upon treatment with dithionite, the NBD-PC on the outer leaflet is converted to non-fluorescent ABD-PC, while the inner leaflet remains unaltered due to the impermeability of dithionite, resulting in approximately a 50% reduction in overall fluores-cence. With the activation of scramblase, there is facilitated translocation of NBD-PC between leaflets, leading to a further diminution of fluorescence intensity over time. NBD-PC, 7-nitro-benz-2-oxa-1,3-diazol-4-yl -PC. ABD-PC, 7-amino-2,1,3-benzoxadiazol-4-yl-PC. b. CLCC1 alone lacks detectable phospholipids scramblase activity *in vitro*. Protein-free liposomes and CLCC1 liposomes were prepared for *in vitro* lipid scrambling, and CLCC1-liposomes were indistin-guishable from protein-free liposomes. c. IB analysis of TMEM41B protein level in CLCC1-TMEM41B complex-liposomes and 0.6 nM to 0.15 nM TMEM41B-liposomes. d. Schematic diagram of liposome-based leakiness assay. During the assembly process, NBD-glucose is encapsulated within liposomes. If the liposome remains intact, adding dithionite to the external buffer would not quench the NBD-glucose signal, as dithionite cannot permeate the membrane. However, in compromised liposomes, dithionite penetrates and interacts with NBD-glucose, result-ing in fluorescence quenching. e. NBD-Glucose leakiness assay demonstrating the intactness of protein-free liposomes, 0.3 nM TMEM41B-liposomes, CLCC1-liposomes and CLCC1-TMEM41B complex liposomes. For b-c and e, representative results of at least 3 biological independent replicates are shown.

**Extended Data Figure 7.**
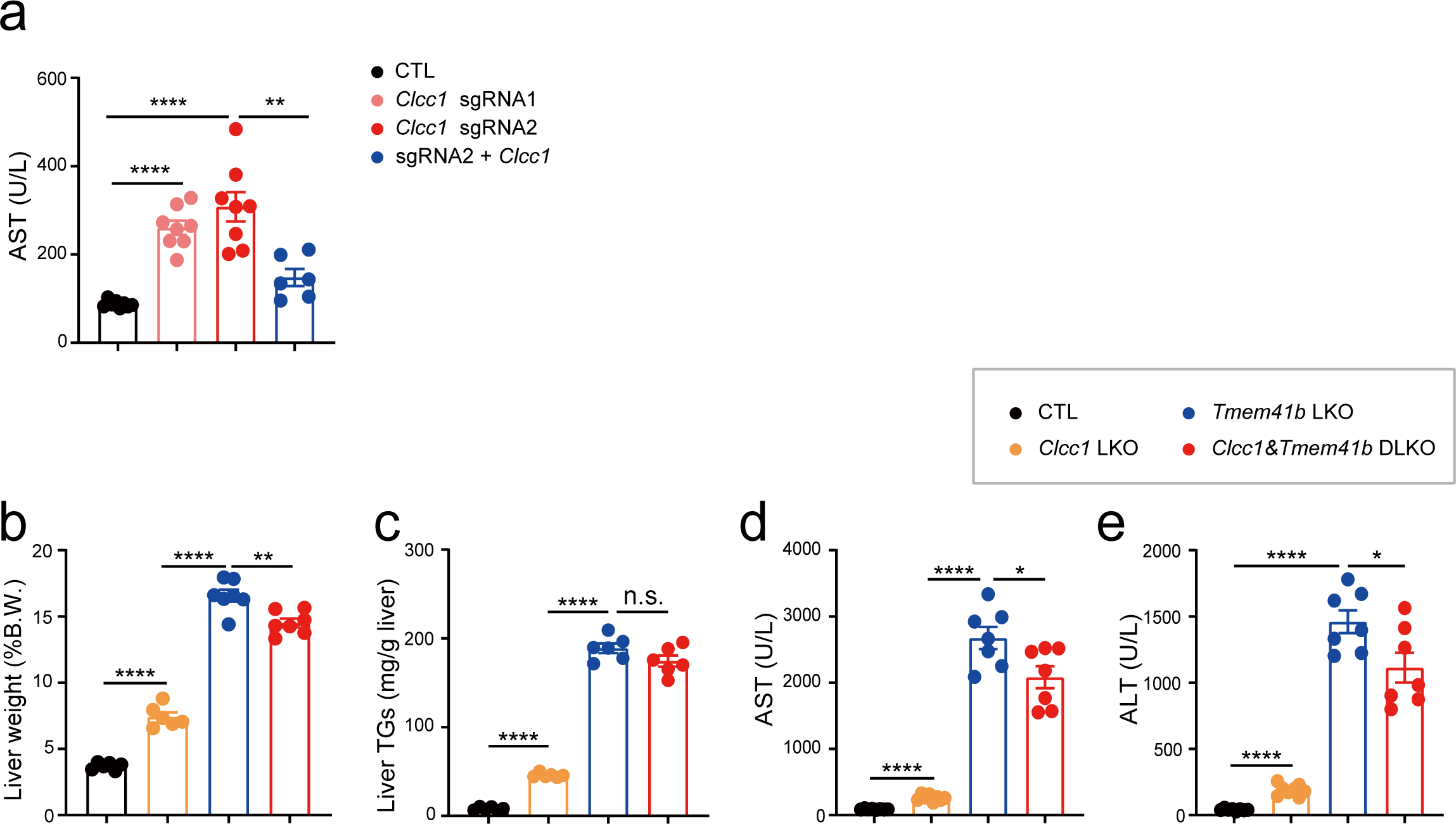
CLCC1 functions with TMEM41B to maintain tissue homeostasis. a. Quantification of plasma AST of mice in Figure 4a. Data are presented as mean ± SEM. **p<0.01, ****p < 0.0001 (two-tailed Student’s t test). b. Quantification of liver weight (% of B.W.) of mice in Figure 4h. Data are presented as mean ± SEM. **p<0.01, ****p < 0.0001 (two-tailed Student’s t test). c. Quantification of liver liver TG levels of mice in Figure 4h. Data are presented as mean ± SEM. n.s., not significant, ****p < 0.0001 (two-tailed Student’s t test). d. Quantification of plasma AST of mice in Figure 4h. Data are presented as mean ± SEM. *p<0.05, ****p < 0.0001 (two-tailed Student’s t test). e. Quantification of plasma ALT of mice in Figure 4h. Data are presented as mean ± SEM. *p<0.05, ****p < 0.0001 (two-tailed Student’s t test).

**Extended Data Figure 8.**
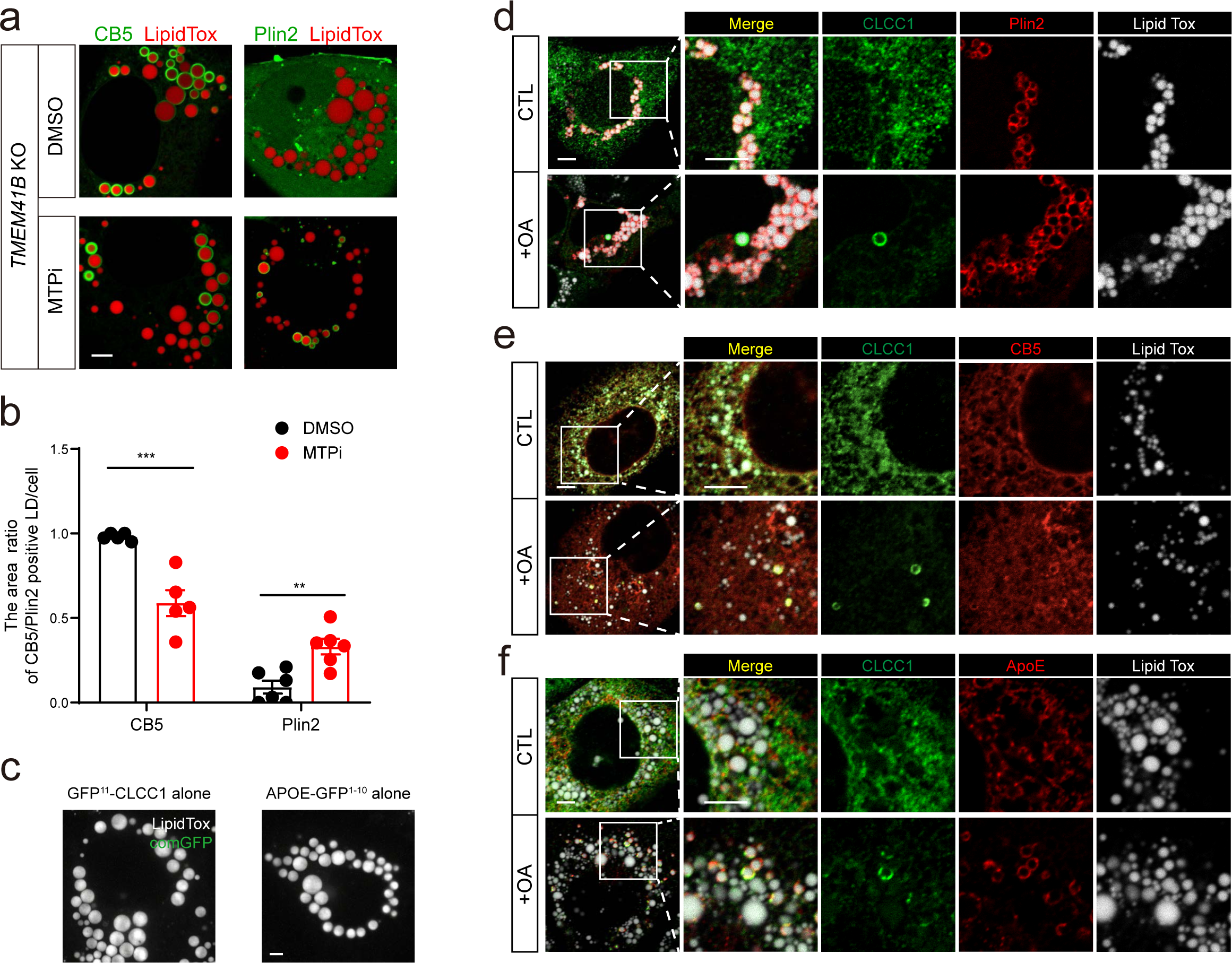
Re-localization of CLCC1 to curved ER with lipid stimuli. a. MTP inhibition redirects neutral lipids from geLD to cLD in TMEM41B deficient cells. CRISPR/-Cas9-mediated *TMEM41B* KO Huh7 cells infected with AAVs expressing GFP-CB5 (left) or GFP-Plin2 (right) were treated with DMSO (upper) or MTP inhibitors (lower) for 24 h. Cells were stained with LipidTox prior to confocal microscopy. Scale bar, 5 μm. b. Quantification of cLD/total LD area from (a). Data are presented as mean ± SEM. **p<0.01, ***p<0.001 (two-tailed Student’s t test). n = 5 cells for CB5 and 6 for Plin2 in both DMSO or MTPi group. c. Neither GFP11-CLCC1 or APOE-GFP1-10 could emit fluorescence in cells. Confocal microscopy of TMEM41B KO cells infected with AAVs expressing either GFP11-CLCC1 or APOE-GFP1-10 alone. Scale bar, 5 μm. d. The absence of Plin2 on CLCC1 encircled LDs induced by OA treatment. WT Huh7 cells infected with AAVs expressing GFP-Plin2, were treated with control (upper) or OA (200 μM) (lower) for 8 h, followed by co-staining with anti-CLCC1 antibody and LipidTox, prior to confocal microscopy anal-ysis. Scale bar, 5 μm. e. OA treatment induced-colocalization of CLCC1 and CB5 surrounding LDs. WT Huh7 cells infected with AAVs expressing GFP-CB5 were treated as in (d). Scale bar, 5 μm. f. OA treatment induced-colocalization of CLCC1 and ApoE surrounding LDs. WT Huh7 cells infected with AAVs expressing ApoE-GFP were treated as in (d). Scale bar, 5 μm. For a and c-f, representative results of at least 3 biological independent replicates are shown.

**Extended Data Figure 9.**
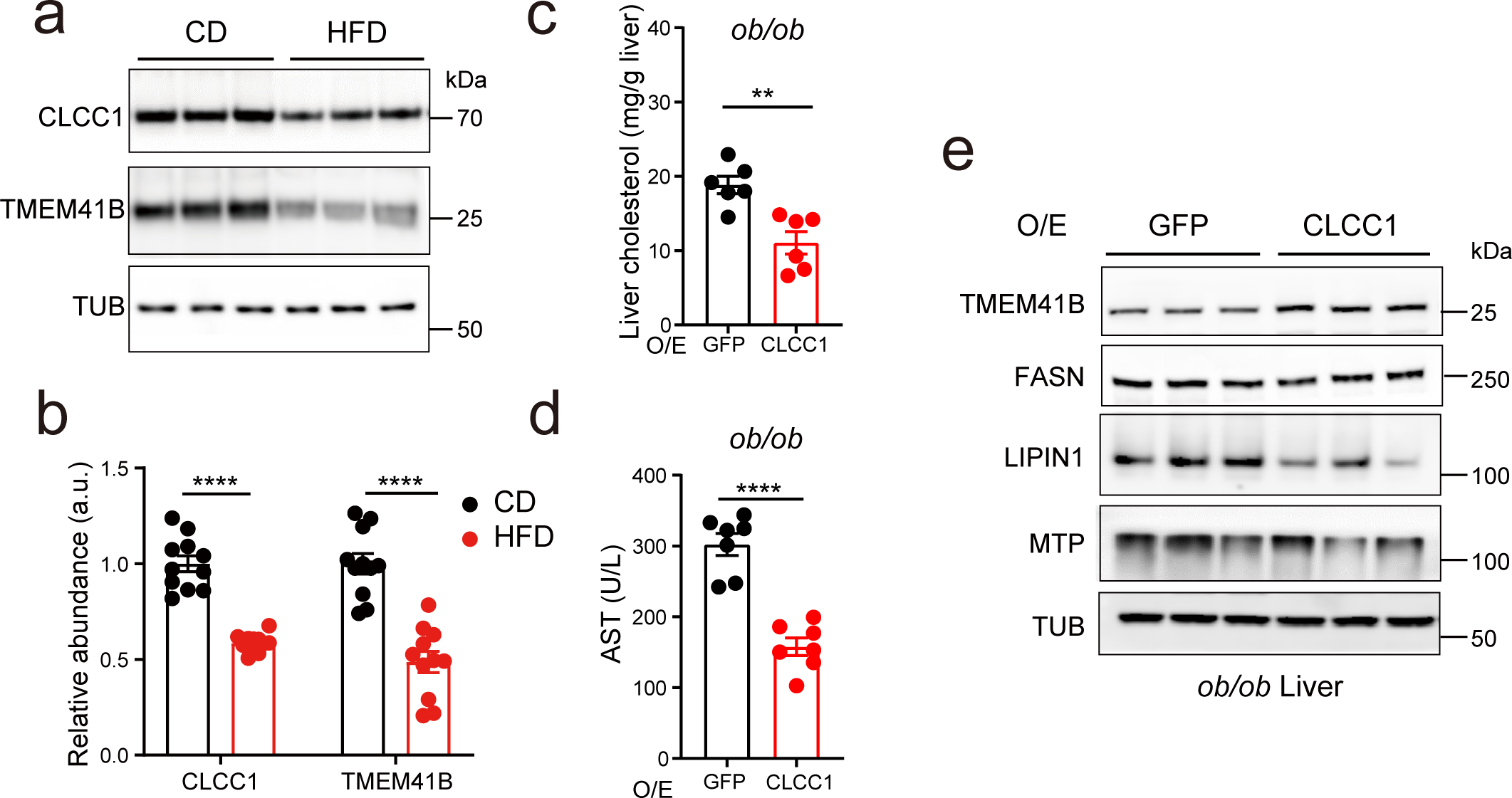
CLCC1 mitigates metabolic dysfunction caused by membrane stress. a. IB analysis of hepatic CLCC1 and TMEM41B protein level in the mice fed with chow diet (CD) or high fat diet (HFD). b. Statistical analysis of IB signal in (a). Data are presented as mean ± SEM. ****p < 0.0001 (two-tailed Student’s t-test). c. Quantification of liver cholesterol levels of mice in Figure 5i. GFP, n = 6; CLCC1, n = 6. Data are presented as mean ± SEM. **p < 0.01 (two-tailed Student’s t test). d. Quantification of plasma AST of mice in Figure 5i. GFP, n = 7; CLCC1, n = 7. Data are presented as mean ± SEM. ****p < 0.0001 (two-tailed Student’s t test). e. IB analysis of liver samples from mice in Figure 5i. For a-b and e, representative results of at least 3 biological independent replicates are shown.

**Extended Data Table 1.**
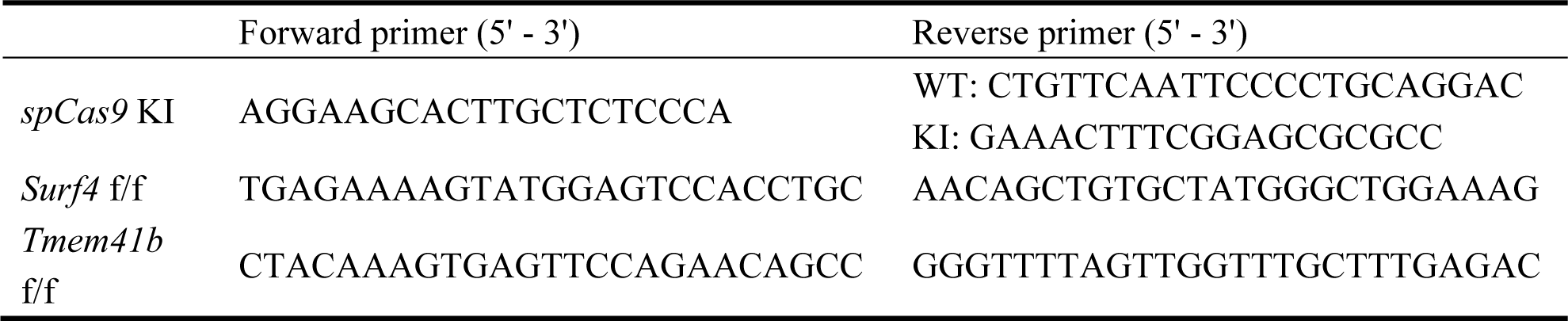
Primer sequences for genotyping.

**Extended Data Table2.**
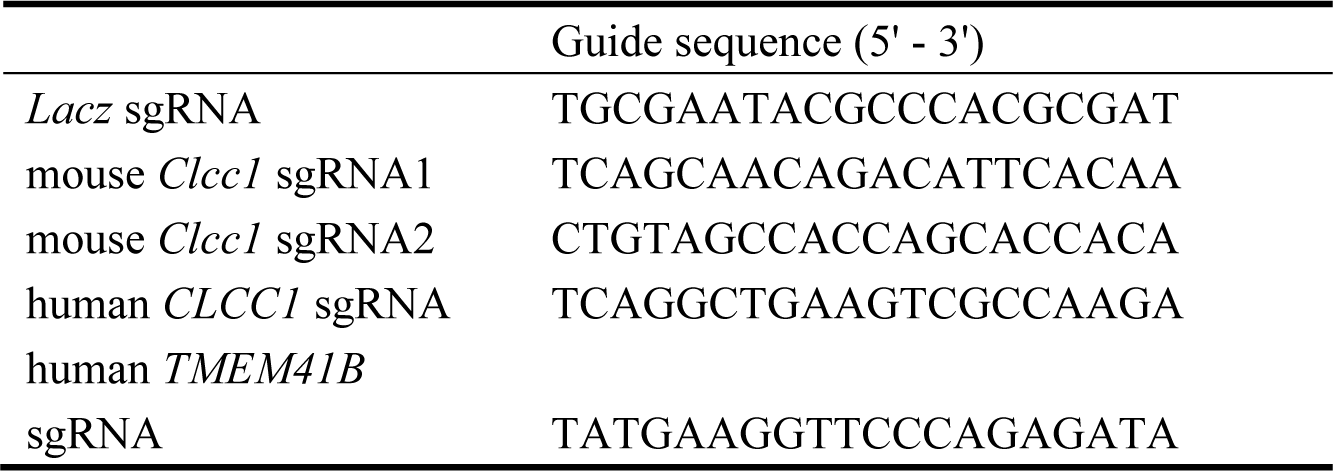
Sequences of oligonucleotides for sgRNA constructs.

## References

1 Mathiowetz, A. J. & Olzmann, J. A. Lipid droplets and cellular lipid flux. Nature Cell Biology, doi:10.1038/s41556-024-01364-4 (2024).

2 Farese, R. V. & Walther, T. The Phase of Fat: Mechanisms and Regulation of Lipid Storage. Biophys J 120, 9a–9a (2021).

3 Collaborators, G. B. D. C. o. D. Global, regional, and national age-sex-specific mortality for 282 causes of death in 195 countries and territories, 1980-2017: a systematic analysis for the Global Burden of Disease Study 2017. Lancet 392, 1736–1788, doi:10.1016/S0140-6736(18)32203-7 (2018).

4 Goldstein, J. L. & Brown, M. S. A century of cholesterol and coronaries: from plaques to genes to statins. Cell 161, 161–172, doi:10.1016/j.cell.2015.01.036 (2015).

5 Gillon, A. D., Latham, C. F. & Miller, E. A. Vesicle-mediated ER export of proteins and lipids. Biochim Biophys Acta 1821, 1040–1049, doi:10.1016/j.bbalip.2012.01.005 (2012).

6 Wang, X. & Chen, X. W. Cargo Receptor-Mediated ER Export in Lipoprotein Secretion and Lipid Homeostasis. Cold Spring Harb Perspect Biol 15, doi:10.1101/cshperspect.a041260 (2023).

7 Sirwi, A. & Hussain, M. M. Lipid transfer proteins in the assembly of apoB-containing lipoproteins. J Lipid Res 59, 1094–1102, doi:10.1194/jlr.R083451 (2018).

8 Fisher, E. A. & Ginsberg, H. N. Complexity in the secretory pathway: The assembly and secretion of apolipoprotein B-containing lipoproteins. Journal of Biological Chemistry 277, 17377–17380, doi:10.1074/jbc.R100068200 (2002).

9 Vance, J. E. Phospholipid synthesis and transport in mammalian cells. Traffic 16, 1–18, doi:10.1111/tra.12230 (2015).

10 Banerjee, S. & Prinz, W. A. Early steps in the birth of four membrane-bound organelles-Peroxisomes, lipid droplets, lipoproteins, and autophagosomes. Curr Opin Cell Biol 84, 102210, doi:10.1016/j.ceb.2023.102210 (2023).

11 Huang, D. et al. TMEM41B acts as an ER scramblase required for lipoprotein biogenesis and lipid homeostasis. Cell Metab 33, 1655–1670 e1658, doi:10.1016/j.cmet.2021.05.006 (2021).

12 Ghanbarpour, A., Valverde, D. P., Melia, T. J. & Reinisch, K. M. A model for a partnership of lipid transfer proteins and scramblases in membrane expansion and organelle biogenesis. Proc Natl Acad Sci U S A 118, doi:10.1073/pnas.2101562118 (2021).

13 Moretti, F. et al. TMEM41B is a novel regulator of autophagy and lipid mobilization. EMBO Rep 19, doi:10.15252/embr.201845889 (2018).

14 Morita, K. et al. Genome-wide CRISPR screen identifies TMEM41B as a gene required for autophagosome formation. Molecular Biology of the Cell 29 (2018).

15 Schneider, W. M. et al. Genome-Scale Identification of SARS-CoV-2 and Pan-coronavirus Host Factor Networks. Cell 184, 120-+, doi:10.1016/j.cell.2020.12.006 (2021).

16 Hoffmann, H. H. et al. TMEM41B Is a Pan-flavivirus Host Factor. Cell 184, 133-+, doi:10.1016/j.cell.2020.12.005 (2021).

17 Nogales, E. & Mahamid, J. Bridging structural and cell biology with cryo-electron microscopy. Nature 628, 47–56, doi:10.1038/s41586-024-07198-2 (2024).

18 Wu, Y. et al. A practical multicellular sample preparation pipeline broadens the application of in situ cryo-electron tomography. J Struct Biol 215, 107971, doi:10.1016/j.jsb.2023.107971 (2023).

19 Xu, C. S. et al. An open-access volume electron microscopy atlas of whole cells and tissues. Nature 599, 147–151, doi:10.1038/s41586-021-03992-4 (2021).

20 Parlakgul, G. et al. Regulation of liver subcellular architecture controls metabolic homeostasis. Nature 603, 736–742, doi:10.1038/s41586-022-04488-5 (2022).

21 Kory, N., Farese, R. V., Jr. & Walther, T. C. Targeting Fat: Mechanisms of Protein Localization to Lipid Droplets. Trends Cell Biol 26, 535–546, doi:10.1016/j.tcb.2016.02.007 (2016).

22 Song, J. et al. Identification of two pathways mediating protein targeting from ER to lipid droplets. Nat Cell Biol 24, 1364–1377, doi:10.1038/s41556-022-00974-0 (2022).

23 Victor, M. B. et al. Lipid accumulation induced by APOE4 impairs microglial surveillance of neuronal-network activity. Cell Stem Cell 29, 1197–1212 e1198, doi:10.1016/j.stem.2022.07.005 (2022).

24 Wang, X. et al. Receptor-Mediated ER Export of Lipoproteins Controls Lipid Homeostasis in Mice and Humans. Cell Metab 33, 350–366 e357, doi:10.1016/j.cmet.2020.10.020 (2021).

25 Graham, S. E. et al. Author Correction: The power of genetic diversity in genome-wide association studies of lipids. Nature 618, E19–E20, doi:10.1038/s41586-023-06194-2 (2023).

26 Platt, R. J. et al. CRISPR-Cas9 knockin mice for genome editing and cancer modeling. Cell, 440-455, doi:10.1016/j.cell.2014.09.014 (2014).

27 Ginsberg, H. N. & Fisher, E. A. The ever-expanding role of degradation in the regulation of apolipoprotein B metabolism. J Lipid Res 50 Suppl, S162-166, doi:10.1194/jlr.R800090-JLR200 (2009).

28 Brodsky, J. L. & Fisher, E. A. The many intersecting pathways underlying apolipoprotein B secretion and degradation. Trends Endocrinol Metab 19, 254–259, doi:10.1016/j.tem.2008.07.002 (2008).

29 Raabe, M. et al. Analysis of the role of microsomal triglyceride transfer protein in the liver of tissue-specific knockout mice. J Clin Invest 103, 1287–1298, doi:10.1172/JCI6576 (1999).

30 Abramson, J. et al. Accurate structure prediction of biomolecular interactions with AlphaFold 3. Nature, doi:10.1038/s41586-024-07487-w (2024).

31 Wu, L., Liu, L., Xu, B., Huang, D. & Chen, X. W. In vitro and in vivo assay of the ER lipid scramblase TMEM41B. STAR Protoc 3, 101333, doi:10.1016/j.xpro.2022.101333 (2022).

32 Ploier, B. & Menon, A. K. A Fluorescence-based Assay of Phospholipid Scramblase Activity. J Vis Exp, doi:10.3791/54635 (2016).

33 Yao, Z., Zhou, H., Figeys, D., Wang, Y. & Sundaram, M. Microsome-associated lumenal lipid droplets in the regulation of lipoprotein secretion. Curr Opin Lipidol 24, 160–170, doi:10.1097/MOL.0b013e32835aebe7 (2013).

34 Wang, Y. W., Tran, K. & Yao, Z. M. The activity of microsomal triglyceride transfer protein is essential for accumulation of triglyceride within microsomes in McA-RH7777 cells - A unified model for the assembly of very low density lipoproteins. Journal of Biological Chemistry 274, 27793–27800, doi:DOI 10.1074/jbc.274.39.27793 (1999).

35 Khatun, I., Walsh, M. T. & Hussain, M. M. Loss of both phospholipid and triglyceride transfer activities of microsomal triglyceride transfer protein in abetalipoproteinemia. J Lipid Res 54, 1541–1549, doi:10.1194/jlr.M031658 (2013).

36 Rader, D. J. & Kastelein, J. J. Lomitapide and mipomersen: two first-in-class drugs for reducing low-density lipoprotein cholesterol in patients with homozygous familial hypercholesterolemia. Circulation 129, 1022–1032, doi:10.1161/CIRCULATIONAHA.113.001292 (2014).

37 Cuchel, M. et al. Efficacy and safety of a microsomal triglyceride transfer protein inhibitor in patients with homozygous familial hypercholesterolaemia: a single-arm, open-label, phase 3 study. Lancet 381, 40–46, doi:10.1016/S0140-6736(12)61731-0 (2013).

38 Guo, L. et al. Disruption of ER ion homeostasis maintained by an ER anion channel CLCC1 contributes to ALS-like pathologies. Cell Res 33, 497–515, doi:10.1038/s41422-023-00798-z (2023).

39 Belloy, M. E., Napolioni, V. & Greicius, M. D. A Quarter Century of APOE and Alzheimer’s Disease: Progress to Date and the Path Forward. Neuron 101, 820–838, doi:10.1016/j.neuron.2019.01.056 (2019).

40 Haney, M. S. et al. APOE4/4 is linked to damaging lipid droplets in Alzheimer’s disease microglia. Nature 628, 154–161, doi:10.1038/s41586-024-07185-7 (2024).

41 Li, L. et al. Mutation in the intracellular chloride channel CLCC1 associated with autosomal recessive retinitis pigmentosa. PLoS Genet 14, e1007504, doi:10.1371/journal.pgen.1007504 (2018).

42 Sanjana, N. E., Shalem, O. & Zhang, F. Improved vectors and genome-wide libraries for CRISPR screening. Nat Methods 11, 783–784, doi:10.1038/nmeth.3047 (2014).

43 Wang, X., Xu, B. L. & Chen, X. W. Acute gene inactivation in the adult mouse liver using the CRISPR-Cas9 technology. STAR Protoc 2, 100611, doi:10.1016/j.xpro.2021.100611 (2021).

44 Ding, Y. et al. Isolating lipid droplets from multiple species. Nat Protoc 8, 43–51, doi:10.1038/nprot.2012.142 (2013).

45 Wittig, I., Braun, H. P. & Schagger, H. Blue native PAGE. Nat Protoc 1, 418–428, doi:10.1038/nprot.2006.62 (2006).

46 Schiotz, O. H. et al. Serial Lift-Out: sampling the molecular anatomy of whole organisms. Nat Methods, doi:10.1038/s41592-023-02113-5 (2023).

47 Mastronarde, D. N. Automated electron microscope tomography using robust prediction of specimen movements. J Struct Biol 152, 36–51, doi:10.1016/j.jsb.2005.07.007 (2005).

48 Turonova, B. et al. Benchmarking tomographic acquisition schemes for high-resolution structural biology. Nat Commun 11, 876, doi:10.1038/s41467-020-14535-2 (2020).

49 Khavnekar, S., Erdmann, P. & Wan, W. TOMOMAN: Streamlining Cryo-electron Tomography and Subtomogram Averaging Workflows Using TOMOgram MANager. Microsc Microanal 29, 1020, doi:10.1093/micmic/ozad067.516 (2023).

50 Zheng, S. Q. et al. MotionCor2: anisotropic correction of beam-induced motion for improved cryo-electron microscopy. Nat Methods 14, 331–332, doi:10.1038/nmeth.4193 (2017).

51 Tegunov, D. & Cramer, P. Real-time cryo-electron microscopy data preprocessing with Warp. Nat Methods 16, 1146–1152, doi:10.1038/s41592-019-0580-y (2019).

52 Konorty, M., Kahana, N., Linaroudis, A., Minsky, A. & Medalia, O. Structural analysis of photosynthetic membranes by cryo-electron tomography of intact Rhodopseudomonas viridis cells. J Struct Biol 161, 393–400, doi:10.1016/j.jsb.2007.09.014 (2008).

53 Meng, E. C. et al. UCSF ChimeraX: Tools for structure building and analysis. Protein Sci 32, e4792, doi:10.1002/pro.4792 (2023).

